# Nuclear pore complexes undergo Nup221 exchange during blood stage asexual replication of *Plasmodium* parasites

**DOI:** 10.1101/2024.02.16.580747

**Authors:** James Blauwkamp, Sushma V Ambekar, Tahir Hussain, Gunnar R Mair, Josh R. Beck, Sabrina Absalon

**Affiliations:** Indiana University School of Medicine, Department of Pharmacology and Toxicology; Iowa State University, Department of Biomedical Sciences; School of Biological Sciences, Queen’s University Belfast

## Abstract

*Plasmodium* parasites, the causative agents of malaria, undergo closed mitosis without breakdown of the nuclear envelope. Unlike closed mitosis in yeast, *P. berghei* parasites undergo multiple rounds of asynchronous nuclear divisions in a shared cytoplasm which results in a multinucleated organism prior to formation of daughter cells within an infected red blood cell. During this replication process, intact nuclear pore complexes (NPCs) and their component nucleoporins play critical roles in parasite growth, facilitating selective bi-directional nucleocytoplasmic transport and genome organization. Here we utilize ultrastructure expansion microscopy (U-ExM) to investigate *P. berghei* nucleoporins at the single nucleus level throughout the 24-hour blood-stage replication cycle. Our findings reveal that these Nups are evenly distributed around the nuclei and organized in a rosette structure previously undescribed around the centriolar plaque, which is responsible for intranuclear microtubule nucleation during mitosis. By adapting the recombination-induced tag exchange (RITE) system to *P. berghei*, we provide evidence of NPC maintenance, demonstrating Nup221 turnover during parasite asexual replication. Our data shed light on the distribution of NPCs and their homeostasis during the blood-stage replication of *P. berghei* parasites.

**Summary Statement:** Employing ultrastructure expansion microscopy and the RITE system, this study unveils Nup221 turnover in nuclear pore complexes of *Plasmodium* parasites, shedding light on their replication mechanisms.

## Introduction

The survival and growth of eukaryotic organisms relies on the movement of macromolecules between the membrane bound nucleus and the cytosol. This nuclear-cytosolic transport is facilitated by nuclear pore complexes (NPCs) that are embedded into the nuclear envelope. Apart from being a channel for selective bidirectional nucleocytoplasmic transport, the NPC plays a key role in chromatin organization, gene regulation, DNA repair, and maintenance of epigenetic memory in many eukaryotes (Lin and Hoelz 2019). Typically, these NPCs are composed of multiple copies of more than 30 different nucleoporins (Nups) per NPC, with eukaryotic NPCs containing around 1000 proteins total (Hoelz, Debler et al. 2011, Grossman, Medalia et al. 2012, Ori, Banterle et al. 2013, Otsuka and Ellenberg 2018, Lin and Hoelz 2019). Nups are widely conserved in most eukaryotes, including yeasts, humans, and various plant species, and have also been identified in parasitic protozoa such as *Trypanosoma brucei* and *Toxoplasma gondii* (Degrasse and Devos 2010, Courjol, Mouveaux et al. 2017). Nups are separated into two distinct categories; structural Nups that form the pore and the intrinsically disordered phenylalanine-glycine repeat (FG) Nups which regulate nucleocytoplasmic transport (Fig. 1a) (Devos, Dokudovskaya et al. 2006, Beck and Hurt 2017). Structurally, the NPCs establish an inner channel called the inner ring, flanked by the outer rings on the cytosolic and nuclear side of the NPC. After the inner channel is constructed the cytoplasmic filament and the nuclear basket Nups attach to their appropriate side of the pore (Devos, Dokudovskaya et al. 2006, Beck and Hurt 2017). The FG Nups, characterized by their FG repeats, occupy the space inside the inner ring and interact with the cargo to facilitate nuclear-cytosolic transport (Strawn, Shen et al. 2004, Devos, Dokudovskaya et al. 2006, Beck and Hurt 2017).

**Fig. 1:**
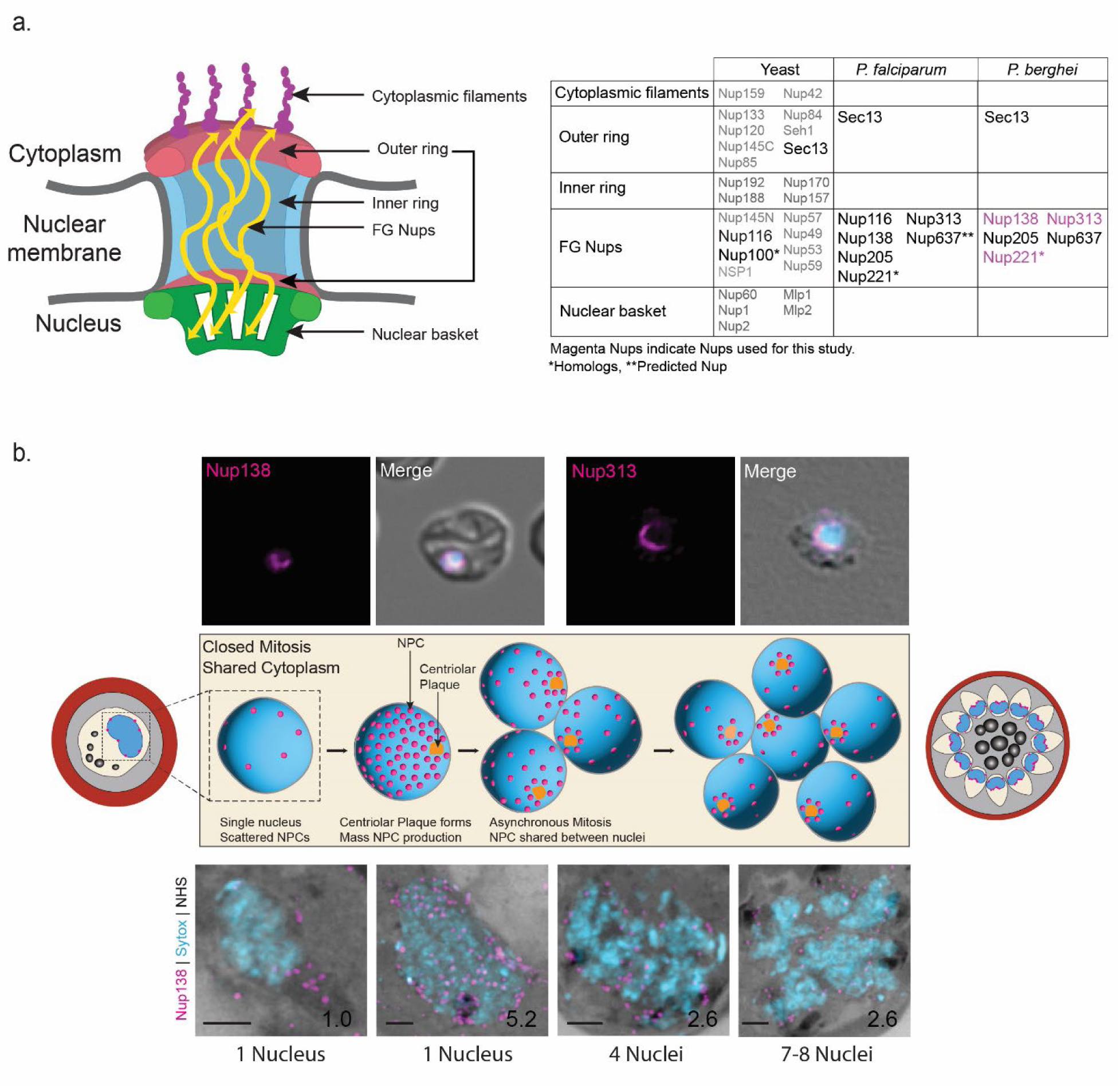
Divergent *P. berghei* NPCs can be analyzed by ultrastructure expansion microscopy (U-ExM). **a.** Schematic representation and table of known nucleoporins (Nups) in yeast, *P. falciparum*, and *P. berghei*. Nups highlighted in magenta indicate Nups visualized in this study, Nups marked with an asterisk represent homologs, Nup637 predicted Nup. **b.** Schematic representation of the *P. berghei* life cycle inside of host red blood cells, showing NPC distribution. Parasites begin with a single nucleus that undergoes mitosis midway through the life cycle. Multiple rounds of asynchronous mitosis lead to a multinucleated parasite with a single cytoplasm before a shared cytokinetic event produces individual parasites. Also shown are comparisons between unexpanded and U-ExM images of Nup138-smHA and Nup313-smHA (magenta) through the life cycle showing differences in these microscopy techniques. Under the U-ExM images lists the number of nuclei in the parasites shown. Scale bars 2 μm, number on images indicates image depth in μm.

One of the most extensively studied organisms with respect to the dynamics and structural aspects of nuclear pore complexes is budding yeast. *Saccharomyces cerevisiae* undergoes closed mitosis during which the nuclear envelope remains intact during karyokinesis. In contrast to open mitosis, where NPC removal during prophase leads to nuclear envelope breakdown, in closed mitosis it has been hypothesized that specific regions of nuclear pore/ complex disassembly cause small breaks in the nuclear envelope, facilitating nuclear envelope fission and formation of two new nuclei (Lénárt, Rabut et al. 2003, Dey, Culley et al. 2020). Like budding yeast, the parasites of the *Plasmodium* genus, the causative agent of malaria, undergo a specialized form of closed mitosis. In these parasites, multiple rounds of asynchronous nuclear division in a shared cytoplasm leads to a multinucleated single-celled parasite (Gerald, Mahajan et al. 2011). While many Nups are conserved through eukaryotes, *Plasmodium* Nups are very divergent, which makes their identification by bioinformatics challenging. In a 2010 study aiming to identify Nups of different eukaryotes, only two Nups were identified in *P. berghei* compared to 19 identified Nups in over 50 different organisms (Neumann, Lundin et al. 2010). Eleven Nups are currently known in the rodent malaria model *P. berghei*; Sec13. Five FG Nups, and five novel Nups identified using Nup313 as bait in proximal labeling assays (Fig. 1a) (Dahan-Pasternak, Nasereddin et al. 2013, Kehrer, Kuss et al. 2018, Ambekar Sushma, Beck Josh et al. 2022). Interestingly, only one of these proteins presents established structural features of eukaryotic Nups (Beck and Hurt 2017).

The limited understanding of essential components of nuclear pore complexes in *Plasmodium* is attributed to the absence of protein homology for Nup identification in *Plasmodium* and the lack of suitable tools to investigate the dynamics of NPCs within relatively small nuclei (a micrometer). While conventional fluorescence and electron microscopy techniques have provided valuable insights into the perinuclear localization of identified Nups in *Plasmodium* parasites, these approaches suffer from low resolution and the significant financial and time investments required for electron microscopy (Doye, Wepf et al. 1994, Winey, Yarar et al. 1997, Weiner, Dahan-Pasternak et al. 2011, Colombi, Webster et al. 2013, Beck and Hurt 2017, Kehrer, Kuss et al. 2018, Dey, Culley et al. 2020, Bertiaux, Balestra et al. 2021). These challenges in comprehending the dynamics and distribution of nuclear pore complexes (NPCs) in *Plasmodium* parasites have spurred the exploration of innovative microscopy techniques to overcome these limitations and gain deeper insights into these vital components. The adaptation of Ultrastructure Expansion Microscopy (U-ExM) to work with parasites now allows for super-resolution imaging of parasites in a cost-effective and high-throughput manner (Gambarotto, Zwettler et al. 2019, Caroline, Charlotta et al. 2021, Liffner and Absalon 2021, Liffner, Cepeda Diaz et al. 2023). U-ExM achieves isotropic expansion of biological samples, increasing their size up to 4.5 times by embedding denatured samples into a hydrogel that expands when exposed to water. Notably, this technique has been employed to achieve sub-nucleus resolution and has successfully visualized Nup313 at the nuclear membrane in *P. falciparum* (Caroline, Charlotta et al. 2021, Liffner, Cepeda Diaz et al. 2023).

In this study, we employed U-ExM to visualize individual NPCs using tagged versions of *P. berghei* Nup138, Nup221, and Nup313. We successfully localized these Nups to the nuclear envelope by employing NHS ester or BodipyFL Ceramide staining throughout the asexual life cycle of the *P. berghei* parasite. Consistent with prior findings using “serial surface” imaging in focused ion beam – scanning electron microscopy (FIB-SEM)(Weiner, Dahan-Pasternak et al. 2011), we observed an increase in NPC number until the parasite’s first mitosis, during which the NPCs were then distributed between the newly developing nuclei. We proceeded to assess NPC assembly and protein turnover using the Recombination Induced Tag Exchange (RITE) system marking the first time this has been accomplished in *Plasmodium*. The RITE system uses a conditional, hormone activated Cre-recombinase to generate permanent epitope tag switch in a target coding sequence. This method has been previously utilized in human cells to observe the dynamic replacement of NPC components in human neurons (Verzijlbergen, Menendez-Benito et al. 2010, Toyama, Arrojo e Drigo et al. 2018).

Leveraging the enhanced resolution of U-ExM coupled with the RITE system, we were able to monitor NPC assembly and maintenance dynamics within a single nucleus during *Plasmodium* replication. In line with previous observations in mammalian cells, we found that NPCs in *P. berghei* undergo protein maintenance after the NPCs have been produced. Remarkably, we also confirmed the presence of Nup221 proteins of different ages in a single NPC by colocalization of original and induced Nup221 signal. Our results expand upon existing data and showcase the new insights achievable through the combination of U-ExM with genetic and biochemical systems.

## Results

### U-ExM visualization of Nups around the nucleus

Previous studies on NPCs in various organisms have predominantly relied on slide-based immunofluorescence assays. For higher resolution, NPCs could be visualized by electron microscopy, but this process is time consuming and costly. The emergence of expansion microscopy in smaller organisms like yeast has significantly advanced NPC visualization and study (Hinterndorfer, Laporte et al. 2022). In our efforts to visualize nucleoporins (Nups) at single NPC resolution in *P. berghei*, we introduced C-terminal endogenous epitope tags on three previously identified *P. berghei* FG Nups: spaghetti monster Hemagglutinin (smHA) tags on Nup138 and Nup313 and a 3xHA/GFP tag on Nup221 (Fig. 2). In comparison to previous studies on unexpanded parasites (Kehrer, Kuss et al. 2018, Ambekar Sushma, Beck Josh et al. 2022), and considering the expected size of the NPCs in yeast (Hinterndorfer, Laporte et al. 2022), expansion microscopy successfully resolves individual NPCs around the nuclear periphery (Fig. 1b). To confirm the position of these Nups at the nuclear membrane, we employed different approaches, visualizing the nuclear membrane using an endoplasmic reticulum (ER) antibody (PfBIP), employing membrane stains such as Bodipy TR Ceramide and Bodipy FL Ceramide, and assessing membrane localization through NHS Ester staining variations around the nucleus (Fig. S1). Among these methods, we determined that differences in NHS ester staining and high concentrations of Bodipy FL Ceramide effectively delineated the nuclear envelope to confidently assign the NPC signal to this structure. In summary, U-ExM allows us to observe individual NPCs on the nuclear envelope.

**Fig. 2:**
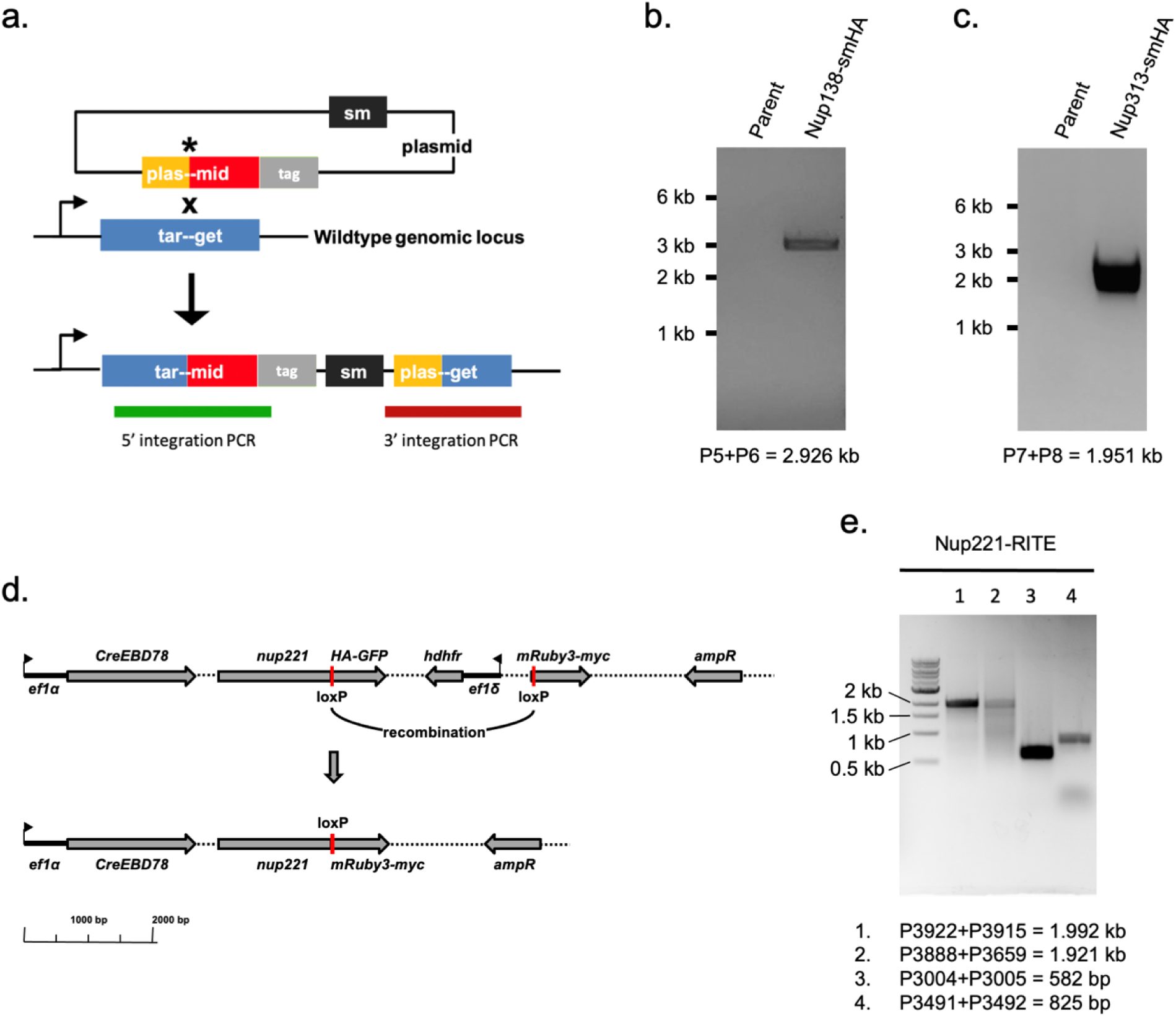
Genotyping of Nup138, Nup313 and Nup221 endogenously tagged parasite lines. **a.** Schematic of general strategy for integration into the endogenous Nup138, Nup313 and Nup221 loci to generate C-terminal tag fusions. The plasmid is linearized with the homology flank targeting each gene (yellow/red box). Following transfection, homologous recombination into the target locus (blue box) results in fusion of the tag to the 3’ end of the gene. Green and red lines show integration PCR products diagnostic for the successful integration. sm, selectable marker. **b,c.** 5’ integration PCRs for Nup138-smHA and Nup313-smHA parasites. **d.** Schematic of the plasmid used to generate the Nup221-RITE parasites. Cre-mediated recombination between loxP sequences removes the 3xHA-GFP tag and hDHFR selection marker and brings the mRuby3-3xMyc tag into frame. **e.** Integration PCRs for the Nup221-RITE parasites. Reactions 1 and 2 are diagnostic for successful 5’ and 3’ integration, respectively. Reactions 3 and 4 are controls amplifying an internal portion of the plasmid and an unrelated genomic locus, respectively.

### *Plasmodium berghei* produces NPCs primarily before mitosis

We next wanted to quantify the number of NPCs surrounding each nucleus during various stages of the parasite’s intraerythrocytic development cycle. To achieve this, we performed U-ExM on parasites that exhibited a range of nuclear numbers and were stained for either Nup138::smHA or Nup313::smHA. In the early stages of the parasite’s life cycle, before the parasites begin their mitotic phase (1n, Ring), we observed Nup138 being uniformly distributed around the nucleus (Fig. 3a), while the Nup313 signal is concentrated at either end of the nucleus (Fig. S2). At this stage, we noticed a higher number of Nup313 foci (approximately 40-50 NPCs) compared to Nup138 (around 25-35 NPCs) (Fig. 3b). As the parasites began mitosis (1n, Trophozoite premitotic and mitotic), there was a significant 5-fold increase in both Nup138 and Nup313 numbers. The parasites produce over 250 NPCs containing Nup313 and over 150 NPCs containing Nup138, distributed around the single nucleus (Fig. 3b). As more nuclei are formed (2-4n Schizont, 5-9n Schizont, 10+n Cytokinesis), the number of NPCs around each nucleus decreased. Initially, approximately 250 NPCs surrounded a single nucleus, which then reduced to around 90-100 NPCs around each of the 2 or 3 nuclei. Eventually, the count further decreases to approximately 25-30 NPCs around 10-12 nuclei (Fig. 3b). Our data suggests that NPC production peaks before mitosis and NPCs are then distributed during each karyokinesis.

**Fig. 3:**
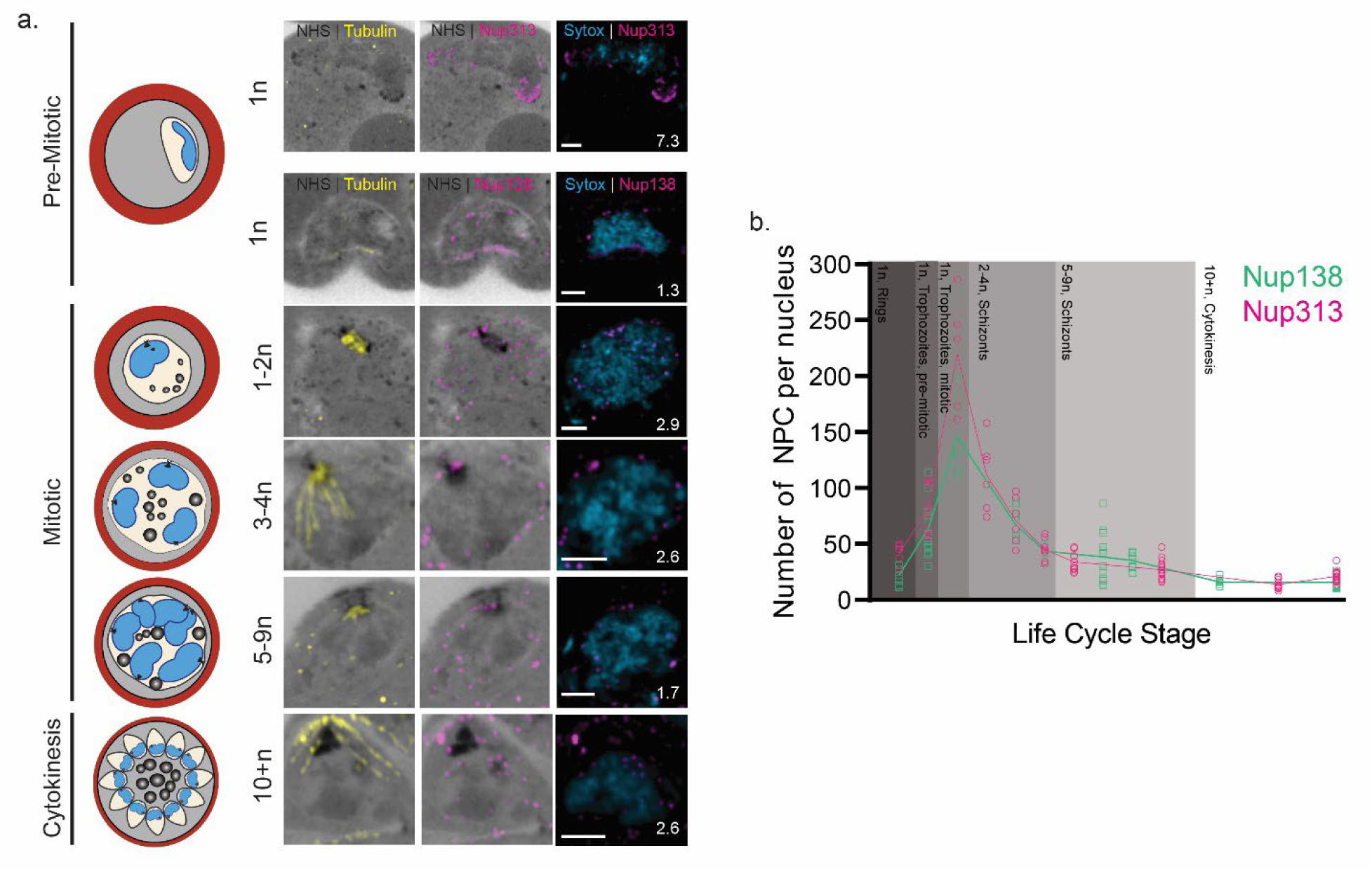
*P. berghei* produces NPCs until mitosis, distributing them among replicated nuclei. **a.** U-ExM images showing Nup313-smHA at early stages and Nup138-smHA through the life cycle of *P. berghei*. Microtubules shown in yellow, Nup138-smHA shown in magenta, DNA shown in blue, and protein density shown in grayscale. Scale bars 2 μm, image depth in μm indicated. **b.** Number of NPCs per nucleus through the *P. berghei* life cycle calculated by counting Nup138-smHA and Nup313 signal. Green indicates Nup138 signal and magenta indicates Nup313-smHA signal.

### *Plasmodium berghei* NPCs are organized around the centriolar plaque

During and after the first mitosis we observe a uniform distribution of NPCs around individual nuclei, as visualized by Nup138 and Nup313 (Fig. 3). Interestingly, upon the first round of mitosis we detect 4-10 foci of Nup138, Nup313, and Nup221, the homolog to yeast Nup100, localizing around the centriolar plaque (CP), which is the nuclear microtubule organization center responsible for intranuclear microtubule nucleation (Fig. 4a)(Arnot, Ronander et al. 2011, Simon, Funaya et al. 2021, Liffner, Cepeda Diaz et al. 2023). Quantification of NPCs around the centriolar plaque reveals a higher count of Nup313 compared to the other two FG Nups used in this study (Fig. 4b). It is important to highlight that after parasite cytokinesis, the centriolar plaque disassembles (Liffner, Cepeda Diaz et al. 2023), but NPCs retain a rosette formation in the vicinity of the former plaque location (Fig. 4c).

**Fig. 4:**
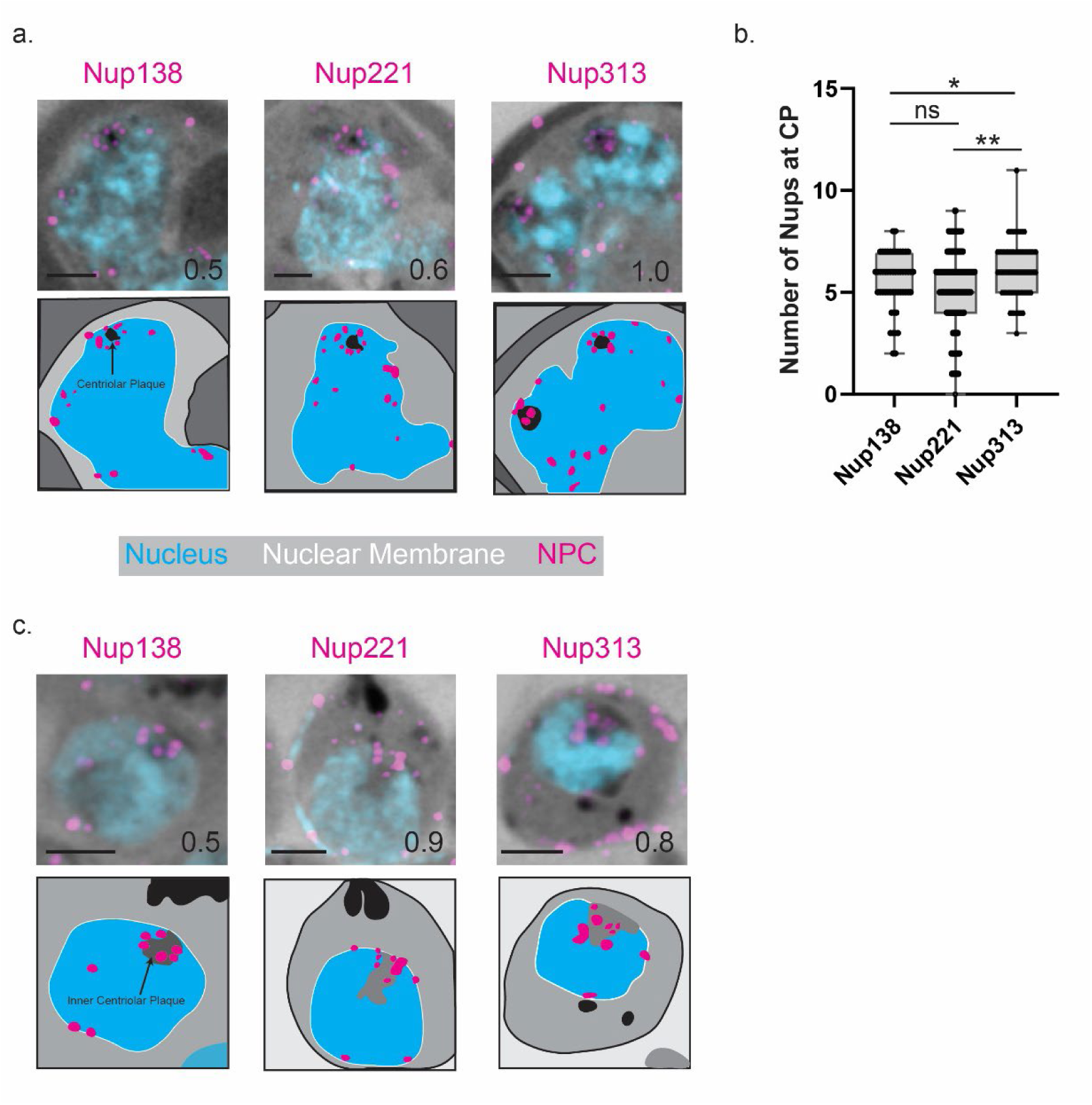
P. berghei NPCs organize around the centriolar plaque (CP) during and after mitosis. **a.** Representative images of mitotic nuclei containing CPs (indicated by black staining of NHS ester) and the organization of three respective FG Nups around the CP. Nups are shown in magenta, DNA is shown in blue, protein density is shown in grayscale. **b.** Quantitation of Nups around the CPs in Nup138-smHA, Nup221-3xHA/GFP, and Nup313-smHA stained NPCs. * indicates P-value > 0.05, ** indicates P-value >0.005. **c.** Representative images of post-mitotic CPs and the organization of three respective FG Nups around where the inner CP disassembles. Nups are shown in magenta, DNA is shown in blue, protein density is shown in grayscale Scale bars 2 μm, image depth in μm indicated on the image.

### The RITE system reveals NPC assembly and maintenance

NPCs in yeast undergo degradation and replacement of Nups and full NPCs (Allegretti, Zimmerli et al. 2020, Lee, Wilfling et al. 2020, Tomioka, Kotani et al. 2020), and that Nup replacement occurs in non-dividing human cells (Toyama, Arrojo e Drigo et al. 2018). Therefore, in order to assess the turnover of Nups in *Plasmodium*, we adapted the Recombination Induced Tag Exchange (RITE) system for the first time in these parasites. The RITE system uses a constitutively expressed Cre recombinase fused to the human estrogen binding domain (EBD) which retains the protein in the cytoplasm by association with HSP70. Upon the addition of β-estradiol, Cre-EBD translocates to the nucleus where it triggers a permanent genetic switch through a recombination event between two *LoxP* flanking the initial tag, removing this tag and bringing a distinct tag into frame (Verzijlbergen, Menendez-Benito et al. 2010) (Fig. S3a). Here we tagged *P. berghei* Nup221 with a RITE system cassette, which includes a 3xHA/GFP tag, a 3xMyc/RFP tag, and the two *LoxP* sites for tag switching (Fig. 2a, 5a). We found that adding as little as 2 nM β-estradiol resulted in a partial switch from HA to Myc tag, while 200 nM β-estradiol induces sufficient relocation into the nucleus for a complete switch from HA to Myc tag in as little as 2 hours, confirmed by PCR analysis (Fig. S3b, c).

**Fig. 5:**
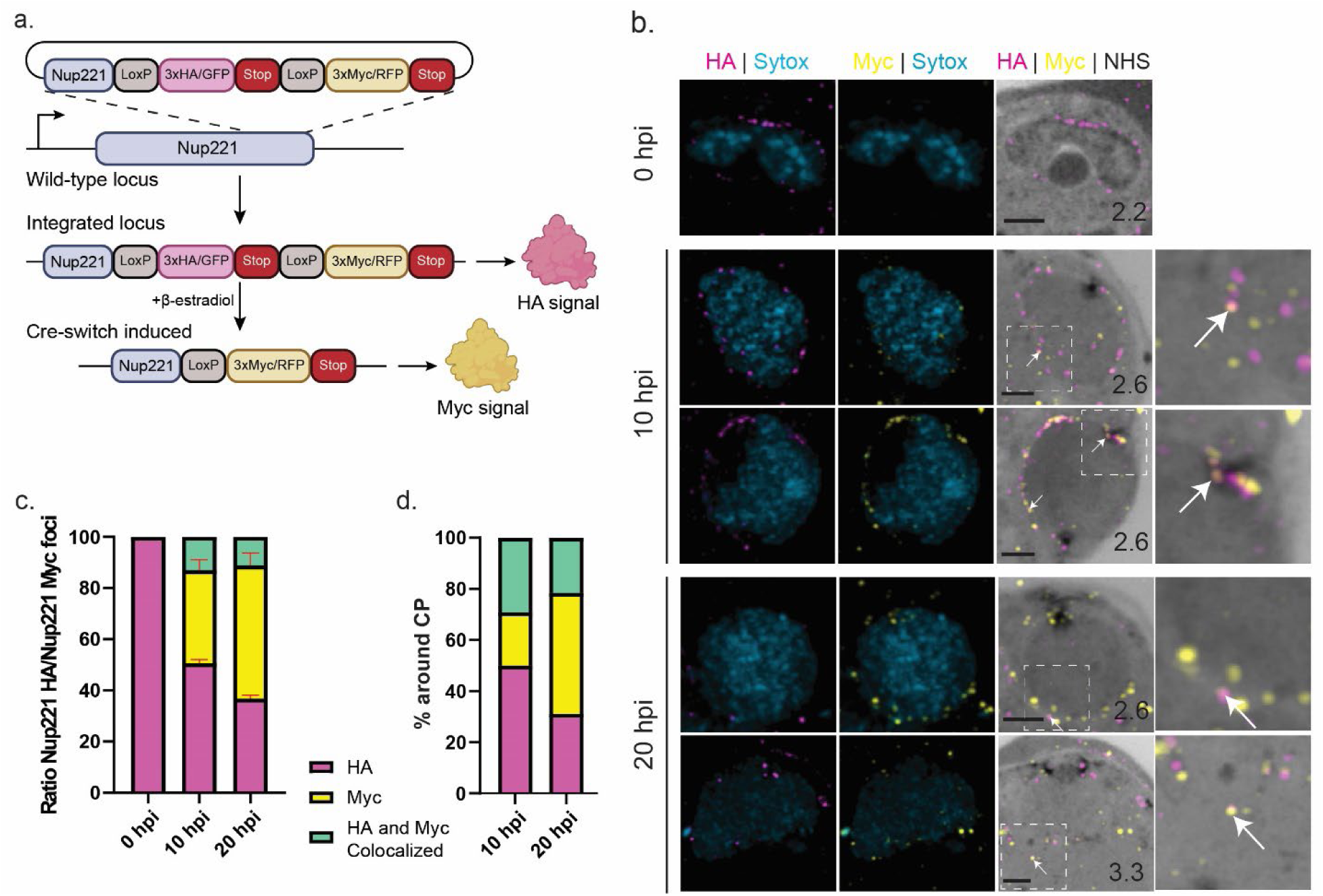
*P. berghei* replaces Nups in post-first mitotic NPCs. **a.** Schematic showing endogenous tagging strategy to introduce the RITE system at the 3’ end of the *Nup221* gene. Translocation of Cre-EBD to the nucleus following treatment with β-estradiol results in excision between loxP sites, removing the original 3xHA/GFP tag while bringing the 3xMyc/RFP tag into frame. **b.** Representative images of *P. berghei* parasites at 0 hours post, 10 hours post, and 20 hours post induction with β-estradiol. HA signal (Scale bars 2 μm, Image depth in μm indicated. White arrows indicate areas of colocalization between HA and Myc. **c.** Ratio Nup221-HA/Nup221-Myc foci in 0 hours-, 10 hours- and 20 hours post induction with β-estradiol. Green bars indicate number of foci that had both HA and Myc signal. **d.** Amount of HA or Myc signal found around the CP in 10 hour- and 20 hour post induction parasites determined by counting foci around the CP. Green bars indicate foci containing both HA and Myc signal.

Initial immunofluorescence imaging over the course of a 16-hour period for both control and 200 nM β-estradiol treated parasites showed visible tag switching by eight hours post induction with β-estradiol (Fig. S4a). By the end of the 16-hour period we observed individual parasites (merozoites) that expressed a mixture of Nup221 tagged with GFP or RFP, both in a typical polar localization. By PCR genotyping, we can detect tag exchange within two hours of estrogen addition. Notably, the continued detection of Nup221-HA through 16 hours post induction indicates a stable population of Nup221 persists for at least eight hours during the parasite life cycle (Fig. S4). To validate complete turnover of Nup221 and ensure no spurious excision was occurring, samples from control and induced conditions were allowed to propagate for an additional 96 hours (four parasite development cycles). While Myc-tagged Nup221 was never detected in control samples, HA-tagged Nup221 was no longer detectable in β-estradiol treated parasites after 96 hours. These results demonstrate that the RITE system is an effective tool for monitoring Nup221 turnover in *P. berghei.* Pairing this technique with U-ExM will provide better resolution for determining NPC maintenance in the parasites.

### *Plasmodium berghei* replaces Nups in post-mitotic NPCs

To investigate NPC assembly and maintenance during and after mitosis, we treated newly invaded parasites (1N) with 200 nM β-estradiol and cultured for 20 hours. We then collected processed samples for U-ExM at 0, 10, and 20 hours post induction with β-estradiol to determine Nup221 presence in NPCs before mitosis occurs (10 hours post induction) and after mitosis, during cytokinesis (20 hours post induction). We hypothesized that if Nup221 is being replaced in the post-mitotic NPCs, we would observe a signal shift from the original HA signal (pink) to the induced Myc signal (yellow), with a higher proportion of induced signal in the 20 hours post induction samples.

At 0 hours post induction, early-stage parasites display only the original HA signal (Fig. 5b). At 10 hours post induction, parasites mature and undergo the first rounds of mitosis resulting in the formation of 2-3 nuclei. These parasites exhibit a mix of the HA and Myc signals, with 50% of foci containing only HA signal, 35% of foci containing only Myc signal, and 15% of foci containing both the HA and Myc signals (Fig. 5b, c). The presence of co-localized HA and Myc signals suggests multiple copies of Nup221 in individual NPCs. By 20 hours post induction with β-estradiol the parasites have stopped producing new NPCs. At this time we found 35% of NPCs contain only HA signal, while 50% of NPCs contain only Myc signal and 15% of NPCs contain both HA and Myc signal (Fig. 5b, c). This result suggests novel Nup221-Myc incorporation into existing NPCs after the parasites have stopped producing them.

Subsequently, we wondered whether the Nups in the NPCs around the CP were subjected to maintenance at the same rate as those around the nuclei. To test this, we specifically monitored the turnover of Nup221 within the NPCs surrounding the CP. At 0 hours post induction, when no CPs are visible, Nup221 is evenly distributed around the parasite nucleus. By 10 hours post induction the CP is present, and now 50% of the NPCs surrounding it contained only the original signal, mirroring observations around the rest of the nucleus. Around 20% of the NPCs contain only the induced signal, a decrease of 15% compared to around the ones around the nucleus, and 30% of the NPCs contain both the original and induced signal, an increase of 15% compared to those scattered around the nucleus (Fig. 5d). This suggests that the turnover of the original Nup221 signal occurs at a slower pace in the NPCs surrounding the nucleus compared to those scattered around it. This trend persists at 20 hours post induction, where 30% of NPCs showed only the original signal, similar to what was seen around the rest of the nucleus. However, 45% of the NPCs contain only the induced signal (a decrease from 50%) and 25% of the NPCs contain both the original and induced signal (an increase from 15%) (Fig. 5d). Altogether, our data indicate that proteins in the NPCs surrounding the CP exhibit greater stability than those in NPCs scattered around the nucleus.

## Discussion

In this study, we applied U-ExM and the RITE System to investigate the distribution and homeostasis of NPCs in the asexual blood stages of the rodent malaria model, *P. berghei*. Previous work on Nups in *Plasmodium* presented challenges due to both the divergent nature of the Nups and the relatively small size of the parasite nucleus at approximately 1 µm. Our results demonstrate the effectiveness of U-ExM in resolving individual NPCs around the parasite nucleus, enabling us to address questions about their individual characteristics. Additionally, for the first time we adapted the RITE system in *Plasmodium,* allowing us to dissect the mechanisms of NPC maintenance. While the RITE system was employed here to monitor protein turnover by tag exchange, it should also be amenable to other conditional mutagenesis strategies, providing an alternative conditional excision strategy in addition to the established dimerizable Cre (DiCre) system (Kent, Modrzynska et al. 2018, Fernandes, Briquet et al. 2020), expanding the genetic toolkit in this important rodent malaria model.

This study defined the number of Nups surrounding the nucleus during the asexual life cycle of *P. berghei*, unraveling a dual localization of NPCs around individual nuclei. Interestingly, we observed a rosette structure around the centriolar plaque (CP) during nuclear replication and this NPC organization persisted in merozoites even when the CP was no longer visible. Our findings align with Ferreira et al, using CryoEM to examine microtubules in *P. falciparum* parasites, revealing a similar ordered NPC structure around the CP during mitosis (Ferreira, Pražák et al. 2023). This circular organization, previously unreported and not observed outside of *Plasmodium*, could suggest a specific role of these NPCs in CP formation, duplication, maintenance, and function during mitosis. For example, *S. cerevisiae* NPC protein Ndc1, a subunit of the transmembrane ring, is a key protein in inserting the nuclear MTOC into the nuclear envelope (Chial, Rout et al. 1998, Jaspersen and Ghosh 2012).

By quantifying individual Nup foci around the nuclei, we determined the number of NPCs per nucleus throughout the life cycle. Our results indicate a rapid increase in NPCs just before the first round of mitosis, consistent with previous observations *in P. falciparum* (Weiner, Dahan-Pasternak et al. 2011). The higher NPC count observed in our study (250 vs 60) might be attributed to greater sample numbers achievable with U-ExM compared to FIB-SEM (our data was taken from 150+ nuclei from 60+ cells over two biological samples) and improved NPC detection by immunostaining against the endogenously tagged Nups.

Previous research in various cell types has shown that the composition of NPCs at specific stages in the cell cycle can influence cell fate (Kane, Rebensburg et al. 2018, Gomar-Alba and Mendoza 2020). In *Plasmodium* parasites it has been shown that SEC13, a component of the nuclear pore complexes and the COPII coat, show a different localization than FG Nups (Kehrer, Kuss et al. 2018). Our study delved into the heterogeneity of NPCs by examining three FG-Nups: Nup138, Nup221 and Nup313. Notably, Nup313 showed a distinct localization in the early ring stage of the parasite suggesting a potential specialized role during this stage of development. Further investigations into the dynamic localization of different types of Nups would be interesting to assess whether Nup313’s temporal differential localization is unique or shared with other Nups that compose the NPC.

Our data indicate that most NPCs contain a single version of Nup221, whether pre-mitotic (Fig. 5c – Nup221-HA, pink) or post-mitotic (Fig. 5c – Nup221-Myc, yellow). However, some NPCs exhibited both pre- and post-mitotic proteins, suggesting a gradual turnover in response to protein degradation or damage (Fig. 5c). These observations echoed findings using the RITE system by Toyama et al., suggesting an active mechanism for NPC maintenance in mammalian cells (Toyama, Arrojo e Drigo et al. 2018). This could also point towards the octagonal rotational symmetry that has been seen in other organisms (Unwin and Milligan 1982, Hinshaw and Milligan 2003). Interestingly, we note a slight increase in protein stability around the CPs, illustrated by higher colocalization between Nup221-HA and Nup221-Myc proteins than around the rest of the nucleus (Fig. 5d), suggesting a higher longevity of this FG Nup in NPCs rosette compared to those scattered across the nucleus.

With the identification of numerous Nups in the *Plasmodium* genome (Kehrer, Kuss et al. 2018, Ambekar Sushma, Beck Josh et al. 2022), and improved tools for visualizing NPCs, we are now poised to investigate the individual functions of Nups. While the exact roles of specific Nups in the *Plasmodium* parasites remains unknown, studies in other organisms have illuminated the involvement of FG Nups in various cellular processes such as protein synthesis (Pulianmackal, Kanakousaki et al. 2022), NPC localization and composition (Wu, Kasper et al. 2001, Ferreira, Stear et al. 2017), as well as RNA transport (Powers, Forbes et al. 1997, Courjol, Mouveaux et al. 2017). While no studies of specific FG Nup function have been conducted in *Plasmodium* parasites, deletions of Nup138 and Nup221 in genome wide screens and individual knock-out attempts have shown all Nups to be essential (Gomes, Bushell et al. 2015, Bushell, Gomes et al. 2017, Zhang, Wang et al. 2018). Likewise, partial knockdown of the Nup SEC13 in human malaria slows growth of the parasite in the blood (Dahan-Pasternak, Nasereddin et al. 2013), while a C-terminal proline enriched sequence from SEC13 in mouse malaria models is required for parasite growth (Kehrer, Kuss et al. 2018). Thus, it is likely that disruption of Nups can lead to growth delays and impaired cell viability in malaria parasites. Our work, along with prior studies, lays the foundation for future research into these divergent Nups as techniques for efficient knockdown continue to advance in malaria parasites.

One of the hallmarks of nuclear envelope breakdown in mammalian open mitosis is the disassembly of NPCs following mitotic kinase signaling (Linder, Köhler et al. 2017, Dey and Baum 2021). This disassembly is a decisive step where multiple Nups are phosphorylated and sequestered to be used upon nuclear envelope reassembly (Kutay, Jühlen et al. 2021). In contrast, *Plasmodium* parasites, like yeast, undergo closed mitosis, where the nuclear envelope remains intact during nuclear division (Gerald, Mahajan et al. 2011). Recent work in fission yeast has used confocal microscopy (Dey, Culley et al. 2020) and expansion microscopy (Hinterndorfer, Laporte et al. 2022) to look at the distribution and dynamics of NPCs in a closed mitosis model. These studies identified NPCs at the bridge between dividing nuclei before nuclear envelope fission. The authors hypothesize that through currently unknown mechanisms, NPC disassembly in the bridge between dividing nuclei contributes to nuclear envelope disassembly, enabling nuclear division. While we observed instances of dividing nuclei with a cluster of NPCs along this bridge, further investigation is needed to explore this mechanism in *Plasmodium* parasites. Identification and characterization of a cytoplasmic facing Nup will allow for this hypothesis to be studied further, and potentially probe into the mechanisms that determine where NPC disassembly occurs and its link to nuclear fission.

In conclusion, our study provides novel insights into the distribution, homeostasis, and homogeneity of NPC complexes in *P. berghei* throughout the asexual life cycle. Understanding the dynamics of these essential components is crucial, as NPCs play a vital role in parasite survival. The methodologies employed here pave the way for further in-depth investigations into the functions of these divergent complexes, facilitating a comprehensive understanding of their role in *Plasmodium* biology. This study sets the stage for future investigations into the diverse Nups in the *Plasmodium* genome and their individual functions. Disruption of Nups, whether structural or FG Nups, likely impacts cell growth and viability in malaria parasites, as observed in related studies. As techniques for knockdown efficiency improvement continue to evolve, exploring the specific functions of Nups in *Plasmodium* is imminent.

## Methods

### Parasite maintenance

*P. berghei* ANKA clone 2.34 and derivatives were maintained in Swiss Webster mice (Charles River). All experiments involving rodents were reviewed and approved by the Iowa State University Institutional Animal Care and Use Committee.

### Genetic modification of *Plasmodium berghei*

Plasmid cloning was carried out using NEBuilder HIFI DNA assembly (NEB). Primer sequences are given in Table 1. To generate a smHA (Viswanathan, Williams et al. 2015) fusion at the endogenous C terminus of *P. berghei* Nup138 the smHA coding sequence was amplified from plasmid pCAG_smFP_HA(Addgene #59759) using primers P4121/4122 and inserted between BamHI/BsiWI immediately downstream of the *Pbnup138* homology region in the plasmid pLIS0654 (Ambekar Sushma, Beck Josh et al. 2022) resulting in the plasmid pSA026. To generate a smHA fusion at the endogenous C terminus of *P. berghei* Nup313, a homology region corresponding to the 3’ coding sequence of *Pbnup313* up to but not including the stop codon was amplified with primers P4140/4126 from *P. berghei* genomic DNA and inserted between AvrII/BamHI in pSA026, replacing the *Pbnup138* homology region and resulting in the plasmid pSA028.

To generate parasites expressing Cre-EBD with the RITE cassette fused to the endogenous C terminus of *P. berghei* Nup221, the Cre-EBD coding sequence was amplified from the plasmid pTWO40 (Terweij, van Welsem et al. 2013) (Addgene #64769) and the RITE cassette was amplified from the plasmid pKV016, which includes a 3xHA-GFP fusion in the pre-excised state and a 3xMYC-mRFP fusion in the post-excised state (Menendez-Benito, van Deventer et al. 2013) (Addgene #64767). These sequences were assembled together with a *Pbnup221* homology region (Ambekar Sushma, Beck Josh et al. 2022), placing Cre-EBD under the control of *P. berghei* EF1 alpha promoter and the hDHFR selectable marker under the control of the *P. berghei* EF1 delta promoter and positioning the hDHFR cassette between the two loxP-containing RITE tags to facilitate marker removed upon excision. The mRFP sequence in the post-excision RITE fusion was subsequently replaced with mRuby3 to yield the final plasmid pL857.

Plasmids pSA026, pSA028 and pL857 were linearized within the *Pbnup138*, *Pbnup313 or Pbnup221* homology region using HindIII, PacI or ScaI, respectively, and transfected into *P. berghei* as described (Hussain, Linera-Gonzalez et al. 2022). Transfected parasites were injected into naïve mice and selected with 0.07 mg/ml pyrimethamine provided *ad libitum* in drinking water beginning 24 hours post-transfection. Parasites were genotyped to confirm integration at *Pbnup138*, *Pbnup313* or *Pbnup221* using PCR strategies described in Supplementary Figures 2 and 4.

### Live imaging and immunofluorescence assays

For live imaging of parasites at regular intervals throughout the experiment, 100μl parasite culture was spun down and resuspended in PBS containing 1μg/ml Hoechst 33342 to stain nuclei. Following a brief incubation at 37℃, parasites were imaged with a 63x objective on an Axio Observer 7 equipped with an Axiocam 702 mono camera and Zen 2.6 Pro software (Zeiss) using the same exposure times for all images across sample groups and experimental replicates.For immunofluorescence assays on unexpanded cells, thin blood smears were air dried and fixed in ice-cold methanol/acetone (50:50) (for detection of smHA tagged Nup138, Nup313 or for detection of Cre). Fixed slides were blocked with Rockland blocking buffer (Rockland) and probed with the primary and secondary antibodies indicated below., Slides were mounted in Fluoromount-G with DAPI (Invitrogen).

### Ultrastructure expansion microscopy

Ultrastructure expansion microscopy was performed as previously described on fixed samples. Briefly, parasites were fixed in 4% PFA/PBS supplemented with 0.0075% glutaraldehyde by the lab of Dr. Josh Beck at Iowa State University and were sent to the lab of Dr. Sabrina Absalon at Indiana University School of Medicine. 12 mm round coverslips (Fisher Cat. No. NC1129240) were treated with poly-D-lysine for 1 hour at 37 °C, washed three times with MilliQ water, and placed in the wells of a 24-well dish. Fixed samples were resuspended at roughly 1.0 % hematocrit in 1xPBS and 500 µL of parasite sample was added to the wells containing the coated coverslips for 30 minutes at 37 °C. Following parasites settling, coverslips were washed 3 times in 1xPBS before treating with 500 µL 1.4% v/v formaldehyde/2% v/v acylamide (FA/AA) in PBS. Samples were incubated overnight at 37 °C. Gelation, denaturation, staining, and expansion of the gels were performed as previously described (Liffner and Absalon 2021). Stained gels were imaged using a Zeiss LSM900 AxioObserver with an Airyscan 2 detector. Images were taken using a 63x Plan-Apochromat objective lens with a numerical aperature of 1.4.

### RITE experiment setup

Asynchronous Nup221::RITE *P. berghei* parasites were collected and allowed to develop overnight in vitro into terminal schizonts. Schizonts were magnetically purified on a MACS LD column (Miltenyi Biotec) and IV injected into naïve mice to establish a synchronous infection. Infections were allowed to develop for three intraerythrocytic developmental cycles (IDCs) to increase parasitemia, then collected at 72 hpi when parasites were predominantly ring stage and passed through a MACS LD column to remove any trophozoites or schizonts by collecting the flow through.

These synchronized ring-stage parasites were used to establish in vitro cultures in complete RPMI containing 200nM β-estradiol or DMSO (vehicle control). Parasites were collected every two hours for live imaging and genomic DNA isolation. After 24 hours, parasites were washed to remove β-estradiol or DMSO and IV injected into naïve mice to monitor Nup turnover in subsequent IDCs. Mice were not provided with pyrimethamine to avoid killing parasites that had undergone excision, which also removes the hDHFR selection marker.

### Antibodies and stains

The following antibodies were used for widefield IFA analysis: rabbit polyclonal anti-HA (1:500, Thermofisher SG77), rabbit anti-Cre polyclonal (1:500, Abcam ab190177). The following antibodies were used for the preparation of stained gels for expansion microscopy: Rat anti-HA (3F10, 1:25, Roche 12158167001), Mouse IgG1 anti-alpha Tubulin Clone B-5-1-2 (1:500, ThermoFisher 32-2500), Rabbit anti-Myc polyclonal (1:500, ThermoFisher 710007). Goat anti-rabbit Alexa Fluor 594 (1:500, ThermoFisher A11012), Goat anti-rat Alexa Fluor 488 (1:500, ThermoFisher A1106), Goat anti-mouse IgG(H+L) Alexa Fluor 555 (1:500, ThermoFisher A21422), and Goat anti-rabbit Alexa Fluor 555 (1:500, ThermoFisher A32732) were used as secondary antibodies. NHS Ester Alexa Fluor 405 in DMSO (8 µg/mL, ThermoFisher A30000) was used to stain proteins and SYTOX™ Deep Red (1 µM, ThermoFisher S11380) was used as a DNA stain.

### Image analysis

All image analysis was done using 3D Airyscan processing at moderate filter strength on Zen Blue Version 3.1 (Zeiss, Oberkochen, Germany). Images shown in this manuscript are maximum intensity projections of between 5 and 60 z-slices of the entire image, with the number indicated as the total Z-depth of the image (Z depth = 0.13 µm * number of slices taken for projection)

### Statistical analysis

Graphical data and statistical analysis was performed on Graphpad Prism version 9.0.

## Acknowledgements

We thank The Company of Biologists Travelling Fellowship grant JCSTF2202676 (S.V.A) for initial travel funding to begin this project. This work was supported by NIH grant AI139579 (G.R.M. and J.R.B.) and was made possible by an award from the Indiana University School of Medicine (S.A., BRG award #<u>2286272</u>). The content is solely the responsibility of the authors and does not necessarily represent the official views of the Indiana University School of Medicine.

## Author Contributions

**Conceptualization:** J.B., S.V.A., G.R.M., J.R.B., and S.A.; **Methodology:** J.B., S.V.A., G.R.M., J.R.B., and S.A.; **Investigation:** J.B., S.V.A., and T.H.; **Data Analysis:** J.B., S.A. **Writing – Original Draft:** J.B.; **Writing – Review and Editing:** J.B., S.V.A., T.H., G.R.M., J.R.B., and S.A; **Funding Acquisition:** S.V.A., G.R.M., J.R.B., and S.A.; **Supervision:** G.R.M., J.R.B. and S.A.

