## Supplemental figures and tables. for "Nuclear pore complexes undergo Nup221 exchange during blood stage asexual replication of *Plasmodium* parasites"

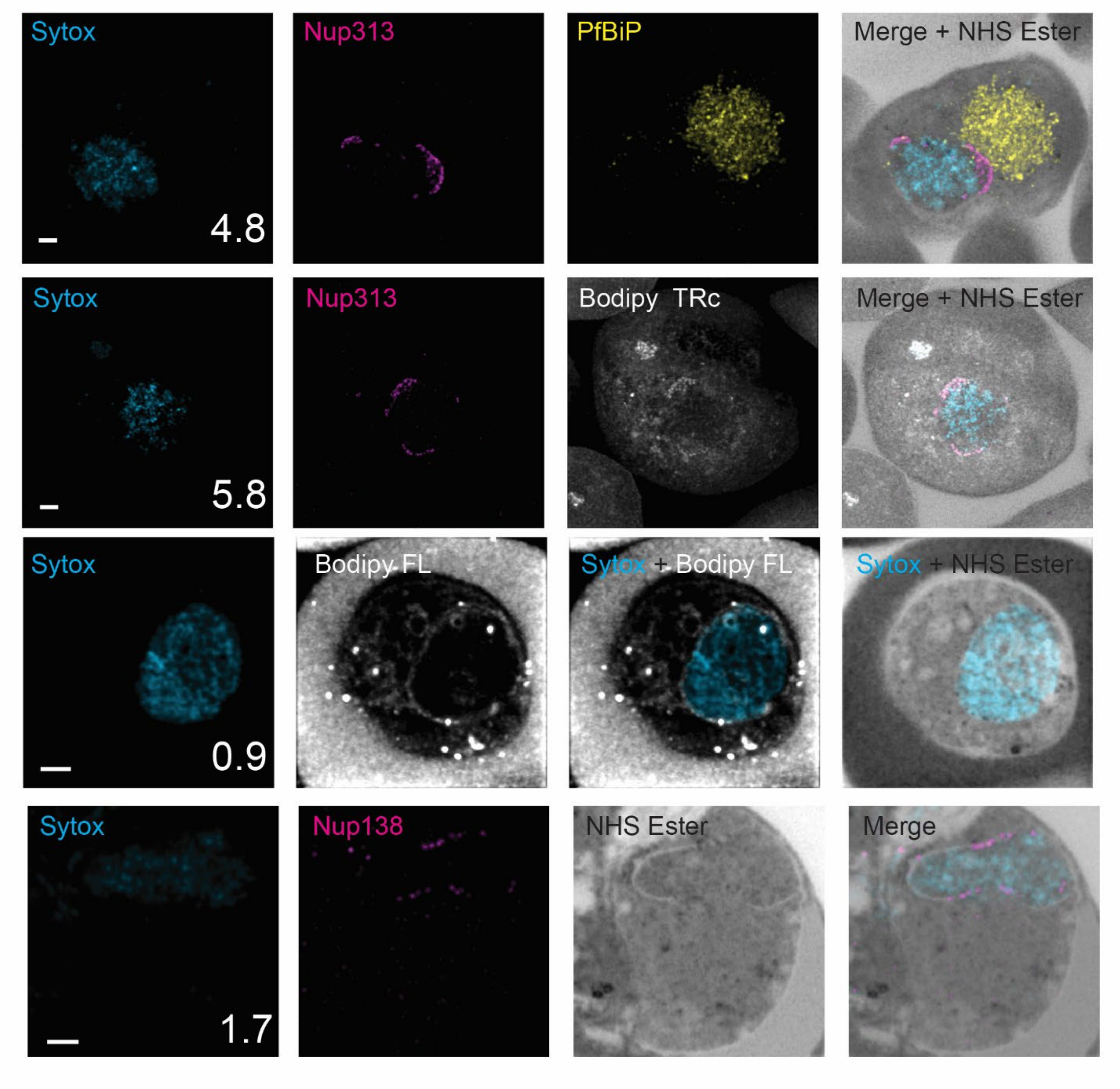


**Fig. S1: Nuclear envelope visualization in *P. berghei*.** Staining with the endoplasmic reticulum protein PfBip (top image, yellow), Bodipy TR ceramide and Bodipy FL ceramide to stain for membranes (middle images, white), and differences in NHS ester staining (grayscale) to visualize the nuclear envelope. Magenta indicates Nup138 or Nup313, scale bars 2 µm, image depth in µm indicated.

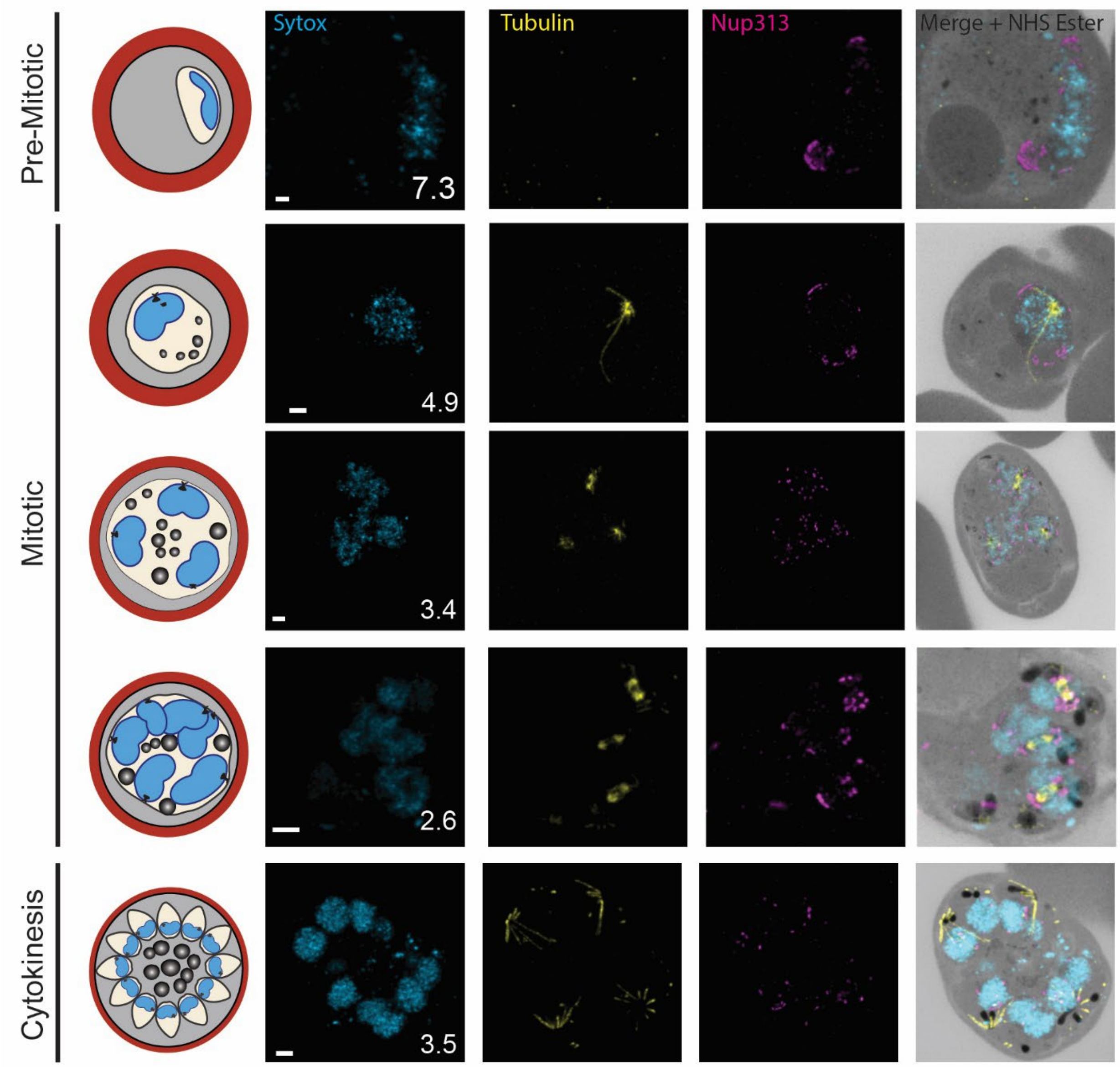


**Fig. S2: Nup313 through the P. berghei blood stage life cycle.** U-ExM images showing Nup313 at early stages and Nup138 through the life cycle of *P. berghei*. DNA shown in blue, microtubules shown in yellow, Nup313 shown in magenta, protein density shown in grayscale. Scale bars 2 µm, image depth in µm indicated.

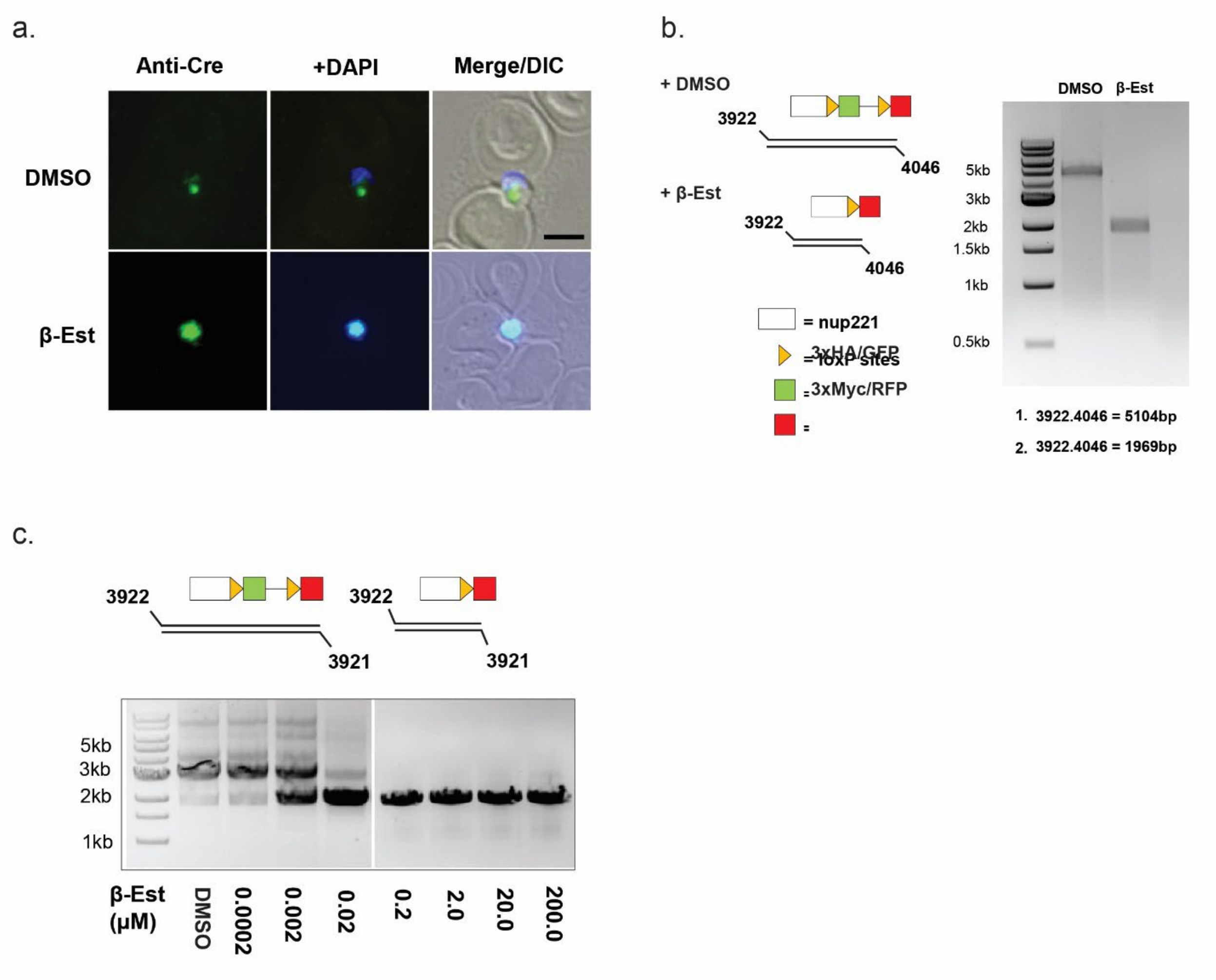


**Fig. S3: Cre-EBD enables inducible genomic excision in *P. berghei.* a.** Immunofluorescence of Cre-EBD expressing *P. berghei* parasites. The addition of β-estradiol to the parasites induces the Cre-EBD to translocate to the nucleus. Scale bar 2 µm. **b.** PCR showing genome shift upon addition of β-estradiol, inducing a tag switch. Primers used are indicated in the diagrams and expected length of PCR products are listed. **c.** PCR showing concentration dependence of the RITE system tested from 0.0002 µM to 200 µM β-estradiol. 2 nM β-estradiol is sufficient to induce a partial tag switch, while 200 nM β-estradiol was sufficient to induce a total tag switch after 2 hours of treatment. PCR primers used are listed in the diagram.

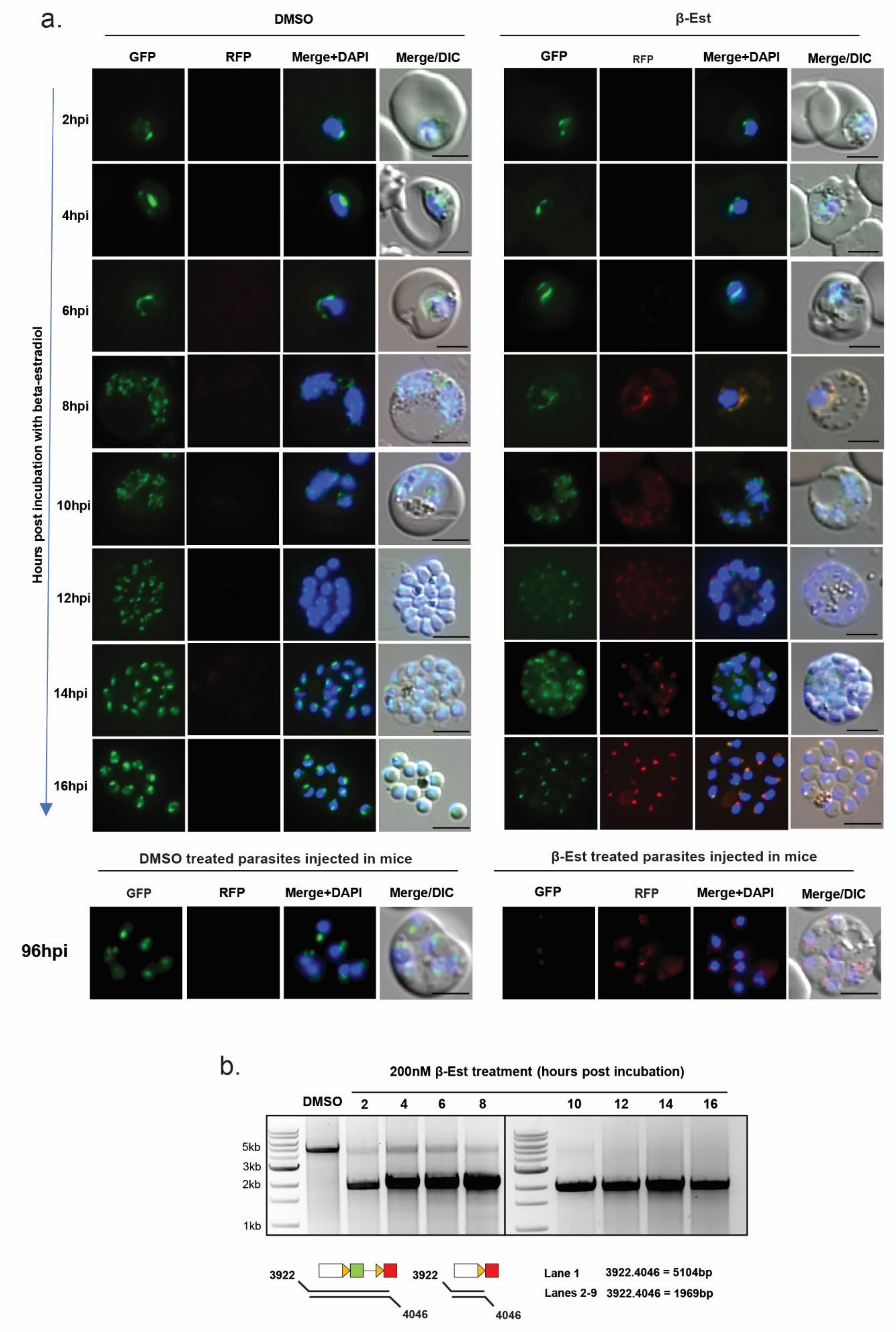
a.
DMSO
β-Est
GFP
RFP
Merge+DAPI
Merge/DIC
Hours post incubation with beta-estradiol
2hpi
4hpi
6hpi
8hpi
10hpi
12hpi
14hpi
16hpi
DMSO treated parasites injected in mice
β-Est treated parasites injected in mice
96hpi
b.
200nM β-Est treatment (hours post incubation)
DMSO
2
4
6
8
10
12
14
16
5kb
3kb
2kb
1kb
3922
4046
Lane 1 3922.4046 = 5104bp
Lanes 2-9 3922.4046 = 1969bp

**Fig. S4: Time course of the RITE system in *P. berghei* of Nup221::RITE. a.** Live imaging of Nup221::RITE parasites showing control or β-estradiol treated parasites during 16 hours of treatment, samples taken every 2 hours. In addition, parasites were cultured an additional 80 hours, where β-estradiol treated parasites only showed RFP signal and control parasites still only showed GFP signal. Scale bars 2 µm. **b.** PCR gel showing the efficiency of Cre-EBD system over 16 hours. PCR primers and expected band sizes are indicated in the diagram.

**Table S1. Sequences of primers used in this study.**

| Name | Sequence |
|---|---|
| P4121 | GTTATTCTCACGAGCTTAGAAATATGAAGGGATCCATGTACCCTTATGATGTGCCCGATTATGC |
| P4122 | GATAGCCAGCGTAGTCCGGGACGTCGTACGGGTAGCCACCCTTGTAGAGTTCGTC |
| P4140 | GACGATAAAAATGAAAATACAAATCCTAGGGAAATTGAAATCGCTTGCACATATTTGAATC |
| P4126 | CATCATAAGGGTACATGGATCCGTTTATGACATTTTGGCTAGTG |
| P5 | GGAAAAAGCTTAACATTTAACTCTTCATTTTTTAAAACGAGCAC |
| P6 | CCGGGACGTCGTACGGG |
| P7 | CTTAGCCCATGCGAATGCATACTATTAGAAC |
| P8 | GTCCATTAACGTCGCCATCCAACTCC |
| P3922 | TTTAAATGATACAAAACACAGACG |
| P3915 | ATACTAGTAGCGTAATCTGGAACG |
| P3888 | TCAATGATTCATAAATAGTTGGACTTG |
| P3659 | ATCAACAAATTATAAATGACAGAAC |
| P3004 | AAAAGATCTATGGTTGGTTCGCTAAACTG |
| P3005 | AAACAATTGTTAATCATTCTTCTCATATAC |
| P3491 | GATAAATTCTCGAGCCCGGGTATCGATTATTTAGAGAATC |
| P3492 | TCTTTACCTCCGCCGGATCCAATATTTTTAATAACATTTC |
| P4046 | TTCCATGACCACCTTCATACGC |
| P3921 | CGCATGAACTCCTTGATGACG |

**Table S2: Plasmids used in this study.**

| Genotype | Plasmid Name |
|---|---|
| nup138::smHA | pSA026 |
| nup313::smHA | pSA028 |
| nup221::RITE | pL857 |

**Table S3: Number of Nup138::smHA foci around nuclei.** Tables showing the number of Nup138::smHA foci around nuclei in different parasites. The number of foci around the centriolar plaque (CP) is also given.

**Table S4: Number of Nup313::smHA foci around nuclei.** Tables showing the number of Nup313::smHA foci around nuclei in different parasites. The number of foci around the centriolar plaque (CP) is also given.

**Table S5: Number of Nupp221:RITE system.** Tables showing the number of either GFP or Myc foci in Nup221::Rite system parasites. Data also counts the number of foci seen around the centriolar plaque (CP) if there is one visible, showing the number of GFP foci, followed by Myc foci, then colocalized GFP and Myc signal. Multiple sets of numbers represent multiple CPs in the same nucleus and the foci around them.

### Table S3

**Experiment** **20220725 Nup138**

| Cell number | Cell Stage | Nucleus number | Number of nups around nucleus | CP (Y or N) | Number of nups around CP | Total nups in cell |
|---|---|---|---|---|---|---|
| 1 | Ring | 1 | 13 | N | --- | 13 |
| 2 | Troph | 1 | 198 | Y | 6, 7 | 198 |
| 3 | Troph | 1 | 75 | N | --- | 75 |
| 4 | Ring | 1 | 19 | N | --- | 19 |
| 5 | Ring | 1 | 47 | N | --- | 47 |
| 6 | Troph | 1 | 74 | N | --- | 74 |
| 7 | Ring | 1 | 11 | N | --- | 11 |
| 7 | Ring | 1 | 22 | N | --- | 22 |
| 7 | Ring | 1 | 31 | N | --- | 31 |
| 7 | Troph | 1 | 49 | N | --- | 49 |
| 7 | Troph | 1 | 100 | N | --- | 100 |
| 7 | Ring | 1 | 31 | N | --- | 21 |
| 9 | Troph | 1 | 59 | N | --- | 59 |
| 10 | Troph | 1 | 40 | N | --- | 40 |
| 11 | Troph | 1 | 110 | Y | 7 | 110 |
| 13 | Schizont | 1 | 30 | Y | 6 | 119 |
| 13 | Schizont | 2 | 19 | Y | 8 | --- |
| 13 | Schizont | 3 | 26 | Y | 5 | --- |
| 13 | Schizont | 4 | 17 | Y | 5 | --- |
| 13 | Schizont | 5 | 14 | Y | 5 | --- |
| 13 | Schizont | 6 | 13 | Y | 5 | --- |
| 14 | Schizont | 1 | 49 | Y | 6, 4 | 174 |
| 14 | Schizont | 2 | 33 | Y | 8, 7 | --- |
| 14 | Schizont | 3 | 50 | Y | 6, 6 | --- |
| 14 | Schizont | 4 | 42 | Y | 7, 5 | --- |
| 15 | Troph | 1 | 117 | Y | 5 | 117 |
| 16 | Troph | 1 | 173 | Y | 6, 7 | 173 |
| 17 | Troph | 1 | 52 | N | --- | 52 |
| 19 | Troph | 1 | 30 | Y | 5 | 30 |
| 19 | Troph | 2 | 42 | Y | 2, 2 | 42 |

**Experiment** **20221121 Nup138 late enriched**

| Cell number | Cell Stage | Nucleus number | Number of nups around nucleus | CP (Y or N) | Number of nups around CP | Total nups in cell |
|---|---|---|---|---|---|---|
| 1 | Merozoites | 1 | 19 | N | --- | 159 |
| 1 | Merozoites | 2 | 15 | N | --- | --- |
| 1 | Merozoites | 3 | 15 | N | --- | --- |
| 1 | Merozoites | 4 | 12 | N | --- | --- |
| 1 | Merozoites | 5 | 22 | N | --- | --- |
| 1 | Merozoites | 6 | 14 | N | --- | --- |
| 1 | Merozoites | 7 | 17 | N | --- | --- |
| 1 | Merozoites | 8 | 14 | N | --- | --- |
| 1 | Merozoites | 9 | 16 | N | --- | --- |
| 1 | Merozoites | 10 | 15 | N | --- | 15 |
| 2 | Ring | 1 | 27 | N | --- | 248 |
| 3 | Schizont | 1 | 29 | Y | 6, 5 | --- |
| 3 | Schizont | 2 | 39 | Y | 6, 6 | --- |
| 3 | Schizont | 3 | 42 | Y | 6, 6 | --- |
| 3 | Schizont | 4 | 43 | Y | 5, 7 | --- |
| 3 | Schizont | 5 | 38 | Y | 7, 7 | --- |
| 3 | Schizont | 6 | 30 | Y | 5, 5 | --- |
| 3 | Schizont | 7 | 24 | Y | 6, 7 | --- |
| 5 | Schizont | 1 | 57 | Y | 3 | 186 |
| 5 | Schizont | 2 | 74 | Y | 6, 7 | --- |
| 5 | Schizont | 3 | 55 | Y | 5 | --- |
| 6 | Schizont | 1 | 62 | Y | 6, 5 | 341 |
| 6 | Schizont | 2 | 60 | Y | 3, 5 | --- |
| 6 | Schizont | 3 | 42 | Y | 6, 7 | --- |
| 6 | Schizont | 4 | 44 | Y | 4, 3 | --- |
| 6 | Schizont | 5 | 86 | Y | 6, 7 | --- |
| 6 | Schizont | 6 | 47 | Y | 3, 4 | --- |
| 7 | Schizont | 1 | 86 | Y | 6, 6 | 216 |
| 7 | Schizont | 2 | 77 | Y | 7 | --- |
| 7 | Schizont | 3 | 53 | Y | 5 | --- |
| 7 | Troph | 1 | 114 | Y | 5, 5 | 114 |
| 8 | Merozoites | 1 | 14 | N | --- | 189 |
| 8 | Merozoites | 2 | 13 | N | --- | --- |
| 8 | Merozoites | 3 | 14 | N | --- | --- |
| 8 | Merozoites | 4 | 23 | N | --- | --- |
| 8 | Merozoites | 5 | 16 | N | --- | --- |
| 8 | Merozoites | 6 | 10 | N | --- | --- |
| 8 | Merozoites | 7 | 12 | N | --- | --- |
| 8 | Merozoites | 8 | 11 | N | --- | --- |
| 8 | Merozoites | 9 | 14 | N | --- | --- |
| 8 | Merozoites | 10 | 14 | N | --- | --- |
| 8 | Merozoites | 11 | 13 | N | --- | --- |
| 8 | Merozoites | 12 | 12 | N | --- | --- |
| 8 | Merozoites | 13 | 13 | N | --- | --- |
| 8 | Merozoites | 14 | 10 | N | --- | --- |
| 9 | Troph | 1 | 131 | Y | 5, 5 | 131 |
| 11 | Merozoites | 1 | 17 | N | --- | 250 |
| 11 | Merozoites | 2 | 15 | N | --- | --- |
| 11 | Merozoites | 3 | 26 | N | --- | --- |
| 11 | Merozoites | 4 | 19 | N | --- | --- |
| 11 | Merozoites | 5 | 19 | N | --- | --- |
| 11 | Merozoites | 6 | 17 | N | --- | --- |
| 11 | Merozoites | 7 | 12 | N | --- | --- |
| 11 | Merozoites | 8 | 20 | N | --- | --- |
| 11 | Merozoites | 9 | 21 | N | --- | --- |
| 11 | Merozoites | 10 | 19 | N | --- | --- |
| 11 | Merozoites | 11 | 18 | N | --- | --- |
| 11 | Merozoites | 12 | 12 | N | --- | --- |
| 11 | Merozoites | 13 | 20 | N | --- | --- |
| 11 | Merozoites | 14 | 15 | N | --- | --- |
| 13 | Troph | 1 | 79 | Y | 6, 7 | 79 |

### Table S4

**Experiment** **20220630 Nup313 berghei**

| Cell number | Cell Stage | Nucleus number | Number of nups around nucleus | CP (Y or N) | Number of nups around CP | Total nups in cell |
|---|---|---|---|---|---|---|
| 1 | Troph | 1 | 50 | N | --- | 50 |
| 2 | Ring | 1 | 27 | N | --- | 27 |
| 2 | Ring | 1 | 63 | N | --- | 63 |
| 3 | Schizont | 1 | 35 | Y | 8, 7 | 191 |
| 3 | Schizont | 2 | 19 | Y | 8 | --- |
| 3 | Schizont | 3 | 16 | Y | 6 | --- |
| 3 | Schizont | 4 | 29 | Y | 8, 6 | --- |
| 3 | Schizont | 5 | 32 | Y | 8,11 | --- |
| 3 | Schizont | 6 | 16 | Y | 7 | --- |
| 3 | Schizont | 7 | 18 | Y | 8 | --- |
| 3 | Schizont | 8 | 26 | Y | 7, 7 | --- |
| 4 | Schizont | 1 | 47 | Y | 6 | 193 |
| 4 | Schizont | 2 | 44 | Y | 3,4 | --- |
| 4 | Schizont | 3 | 59 | Y | 6 | --- |
| 4 | Schizont | 4 | 43 | Y | 4, 5 | --- |
| 6 | Merozoites | 1 | 23 | Y | 5 | 295 |
| 6 | Merozoites | 2 | 25 | Y | 7 | --- |
| 6 | Merozoites | 3 | 24 | Y | 4 | --- |
| 6 | Merozoites | 4 | 35 | Y | 7 | --- |
| 6 | Merozoites | 5 | 25 | Y | 7 | --- |
| 6 | Merozoites | 6 | 18 | Y | 5 | --- |
| 6 | Merozoites | 7 | 16 | Y | 7 | --- |
| 6 | Merozoites | 8 | 21 | Y | 6 | --- |
| 6 | Merozoites | 9 | 21 | Y | 8 | --- |
| 6 | Merozoites | 10 | 20 | Y | 6 | --- |
| 6 | Merozoites | 11 | 17 | Y | 6 | --- |
| 6 | Merozoites | 12 | 14 | Y | 5 | --- |
| 6 | Merozoites | 13 | 20 | Y | 5 | --- |
| 6 | Merozoites | 14 | 16 | Y | 5 | --- |
| 7 | Ring | 1 | 45 | N | --- | 45 |
| 8 | Ring | 1 | 48 | N | --- | 48 |
| 8 | Ring | 1 | 58 | N | --- | 58 |
| 10 | Ring | 1 | 37 | N | --- | 37 |
| 11 | Troph | 1 | 103 | N | --- | 103 |
| 12 | Troph | 1 | 104 | N | --- | 104 |
| 14 | Ring | 1 | 76 | N | --- | 76 |
| 14 | Schizont | 1 | 47 | Y | 6, 6 | 247 |
| 14 | Schizont | 2 | 31 | Y | 6, 7 | --- |
| 14 | Schizont | 3 | 29 | Y | 6, 7 | --- |
| 14 | Schizont | 4 | 19 | Y | 5, 4 | --- |
| 14 | Schizont | 5 | 24 | Y | 4, 5 | --- |
| 14 | Schizont | 6 | 32 | Y | 5, 7 | --- |
| 14 | Schizont | 7 | 38 | Y | 5, 6 | --- |
| 14 | Schizont | 8 | 27 | Y | 6, 7 | --- |
| 15 | Troph | 1 | 83 | N | --- | 83 |
| 16 | Troph | 1 | 246 | N | --- | 246 |
| 16 | Troph | 1 | 173 | N | --- | 173 |
| 17 | Ring | 1 | 57 | N | --- | 57 |
| 18 | Troph | 1 | 233 | Y | 7 | 233 |
| 19 | Ring | 1 | 108 | N | --- | 108 |

Experiment 20221121 Nup313 late enriched

| Cell number | Cell Stage | Nucleus number | Number of nups around nucleus | CP (Y or N) | Number of nups around CP | Total nups in cell |
|---|---|---|---|---|---|---|
| 2 | Schizont | 1 | 56 | Y | 5, 7 | 167 |
| 2 | Schizont | 2 | 45 | Y | 3, 6 | --- |
| 2 | Schizont | 3 | 32 | Y | 6 | --- |
| 2 | Schizont | 4 | 34 | Y | 6 | --- |
| 3 | Troph | 1 | 286 | Y | 6, 7 | 286 |
| 4 | Troph | 1 | 158 | Y | 6 | 283 |
| 4 | Troph | 2 | 125 | Y | 8 | --- |
| 5 | Troph | 1 | 128 | Y | 6 | 231 |
| 5 | Troph | 2 | 103 | Y | 5 | --- |
| 6 | Schizont | 1 | 39 | Y | 5, 5 | 150 |
| 6 | Schizont | 2 | 25 | Y | 4, 5 | --- |
| 6 | Schizont | 3 | 24 | Y | 6, 6 | --- |
| 6 | Schizont | 4 | 28 | Y | 7, 4 | --- |
| 6 | Schizont | 5 | 34 | Y | 5, 6 | --- |
| 7 | Schizont | 1 | 91 | Y | 5, 7 | 198 |
| 7 | Schizont | 2 | 63 | Y | 3, 4 | --- |
| 7 | Schizont | 3 | 44 | Y | 6 | --- |
| 8 | Schizont | 1 | 39 | Y | 4, 5 | 189 |
| 8 | Schizont | 2 | 47 | Y | 5, 5 | --- |
| 8 | Schizont | 3 | 30 | Y | 6, 6 | --- |
| 8 | Schizont | 4 | 27 | Y | 3, 3 | --- |
| 8 | Schizont | 5 | 46 | Y | 3,5 | --- |
| 9 | Troph | 1 | 161 | Y | 2 | 161 |
| 10 | Troph | 1 | 82 | Y | 5, 6 | 156 |
| 10 | Troph | 2 | 74 | Y | 6, 3 | --- |
| 12 | Merozoites | 1 | 18 | N | --- | 164 |
| 12 | Merozoites | 2 | 13 | N | --- | --- |
| 12 | Merozoites | 3 | 23 | N | --- | --- |
| 12 | Merozoites | 4 | 15 | N | --- | --- |
| 12 | Merozoites | 5 | 8 | N | --- | --- |
| 12 | Merozoites | 6 | 13 | N | --- | --- |
| 12 | Merozoites | 7 | 11 | N | --- | --- |
| 12 | Merozoites | 8 | 10 | N | --- | --- |
| 12 | Merozoites | 9 | 12 | N | --- | --- |
| 12 | Merozoites | 10 | 20 | N | --- | --- |
| 12 | Merozoites | 11 | 11 | N | --- | --- |
| 12 | Merozoites | 12 | 10 | N | --- | --- |
| 13 | Schizont | 1 | 97 | Y | 4 | 227 |
| 13 | Schizont | 2 | 77 | Y | 6, 5 | --- |
| 13 | Schizont | 3 | 53 | Y | 5 | --- |

### Table S5

**Experiment** **20220721 Nup221 M1 10hpi**

| Cell number | Cell Stage | Nucleus number | Number of GFP Nups | Number of Myc nups | Number of colocalized Nups | CP (Y or N) | Number of nups around CP (GFP, Myc, colocalized) | Total number of nups in nucleus | Total number nups in cell |
|---|---|---|---|---|---|---|---|---|---|
| 1 | Ring | 1 | 21 | 1 | 0 | Y | 5, 0, 0 | 22 | 22 |
| 2 | Schizont | 1 | 27 | 9 | 3 | N | --- | 39 | 261 |
| 2 | Schizont | 2 | 20 | 10 | 5 | N | --- | 35 | --- |
| 2 | Schizont | 3 | 28 | 14 | 3 | N | --- | 45 | --- |
| 2 | Schizont | 4 | 21 | 8 | 3 | N | --- | 32 | --- |
| 2 | Schizont | 5 | 29 | 12 | 4 | N | --- | 45 | --- |
| 2 | Schizont | 6 | 19 | 8 | 1 | N | --- | 28 | --- |
| 2 | Schizont | 7 | 17 | 8 | 1 | N | --- | 26 | --- |
| 2 | Schizont | 8 | 8 | 2 | 1 | N | --- | 11 | --- |
| 3 | Merozoite | 1 | 13 | 6 | 3 | N | --- | 22 | 22 |
| 4 | Troph | 1 | 48 | 24 | 8 | Y | 0. 0. 0 | 80 | 80 |
| 5 | Troph | 1 | 78 | 25 | 5 | N | --- | 108 | 108 |
| | | | | | | | --- | | --- |
| Totals: | --- | --- | 329 | 127 | 37 | --- | --- | 493 | 493 |

**Experiment** 20220721 Nup221 M1 20hpi

| Cell number | Cell Stage | Nucleus number | Number of GFP Nups | Number of Myc nups | Number of colocalized Nups | CP (Y or N) | Number of nups around CP (GFP, Myc, colocalized) | Total number of nups in nucleus | Total number nups in cell |
|---|---|---|---|---|---|---|---|---|---|
| 1 | Schizont | 1 | 23 | 69 | 10 | Y | 3, 0, 3 | 102 | 248 |
| 1 | Schizont | 2 | 14 | 35 | 8 | Y | 3. 2. 1 | 57 | --- |
| 1 | Schizont | 3 | 19 | 27 | 6 | Y | 2. 4. 0 | 52 | --- |
| 1 | Schizont | 4 | 9 | 22 | 6 | Y | 0. 0. 0 | 37 | --- |
| 2 | Troph | 1 | 31 | 61 | 33 | N | --- | 125 | 125 |
| 3 | Schizont | 1 | 8 | 7 | 6 | Y | 0. 0. 0 | 21 | 188 |
| 3 | Schizont | 2 | 28 | 13 | 7 | Y | 6. 0. 2 | 48 | --- |
| 3 | Schizont | 3 | 23 | 16 | 3 | Y | 4. 1. 1 : 0. 5. 1 | 42 | --- |
| 3 | Schizont | 4 | 21 | 13 | 6 | Y | 5. 0. 2 : 6. 1. 0 | 40 | --- |
| 3 | Schizont | 5 | 19 | 10 | 8 | Y | 2. 0. 4 : 5. 0. 3 | 37 | --- |
| 4 | Merozoite | 1 | 15 | 14 | 0 | N | --- | 29 | 29 |
| 5 | Schizont | 1 | 6 | 14 | 3 | Y | 1. 3. 2 | 23 | 315 |
| 5 | Schizont | 2 | 10 | 15 | 2 | Y | 2. 4. 1 | 27 | --- |
| 5 | Schizont | 3 | 8 | 17 | 4 | Y | 0. 3. 4 | 29 | --- |
| 5 | Schizont | 4 | 6 | 7 | 1 | Y | 1. 2. 1 | 14 | --- |
| 5 | Schizont | 5 | 11 | 10 | 3 | Y | 2. 2. 2 | 24 | --- |
| 5 | Schizont | 6 | 20 | 13 | 4 | Y | 3. 3. 0 | 37 | --- |
| 5 | Schizont | 7 | 9 | 14 | 1 | Y | 1. 1. 1 | 24 | --- |
| 5 | Schizont | 8 | 11 | 13 | 2 | Y | 3. 2. 0 | 26 | --- |
| 5 | Schizont | 9 | 6 | 6 | 4 | Y | 1. 0. 2 | 16 | --- |
| 5 | Schizont | 10 | 8 | 5 | 3 | Y | 2. 1. 2 | 16 | --- |
| 5 | Schizont | 11 | 3 | 3 | 4 | Y | 0. 3. 2 | 10 | --- |
| 5 | Schizont | 12 | 8 | 7 | 5 | Y | 1. 0. 3 | 20 | --- |
| 5 | Schizont | 13 | 7 | 7 | 3 | Y | 2. 1. 1 | 17 | --- |
| 5 | Schizont | 14 | 8 | 8 | 4 | Y | 0. 1. 3 | 20 | --- |
| 5 | Schizont | 15 | 6 | 5 | 1 | Y | 0. 4. 1 | 12 | --- |
| 6 | Troph | 1 | 55 | 45 | 6 | N | --- | 106 | 106 |
| 7 | Merozoite | 1 | 10 | 27 | 5 | N | --- | 42 | 378 |
| 7 | Merozoite | 2 | 17 | 27 | 4 | N | --- | 48 | --- |
| 7 | Merozoite | 3 | 11 | 17 | 5 | N | --- | 33 | --- |
| 7 | Merozoite | 4 | 22 | 24 | 5 | N | --- | 51 | --- |
| 7 | Merozoite | 5 | 20 | 7 | 4 | N | --- | 31 | --- |
| 7 | Merozoite | 6 | 11 | 23 | 2 | N | --- | 36 | --- |
| 7 | Merozoite | 7 | 25 | 23 | 3 | N | --- | 51 | --- |
| 7 | Merozoite | 8 | 9 | 9 | 4 | N | --- | 22 | --- |
| 7 | Merozoite | 9 | 14 | 4 | 3 | N | --- | 21 | --- |
| 7 | Merozoite | 10 | 24 | 18 | 1 | N | --- | 43 | --- |
| 8 | Troph | 1 | 63 | 78 | 52 | N | --- | 193 | 193 |
| 9 | Schizont | 1 | 15 | 64 | 12 | Y | 0. 1. 3 | 91 | 246 |
| 9 | Schizont | 2 | 8 | 28 | 2 | Y | 0. 1. 5 : 0. 4. 1 | 38 | --- |
| 9 | Schizont | 3 | 7 | 18 | 2 | Y | 2. 2. 1 | 27 | --- |
| 9 | Schizont | 4 | 15 | 22 | 2 | Y | 3. 3. 2 | 39 | --- |
| 9 | Schizont | 5 | 26 | 22 | 3 | Y | 2. 1. 1 : 2. 5. 0 | 51 | --- |
| Totals: | | | 689 | 887 | 252 | | | 1828 | 1828 |

**Experiment** 20220722 Nup221 M2 10hpi

| Cell number | Cell Stage | Nucleus number | Number of GFP Nups | Number of Myc nups | Number of colocalized Nups | CP (Y or N) | Number of nups around CP (GFP, Myc, colocalized) | Total number of nups in nucleus | Total number nups in cell |
|---|---|---|---|---|---|---|---|---|---|
| 1 | Troph | 1 | 22 | 39 | 17 | N | --- | 78 | 169 |
| 1 | Troph | 2 | 31 | 43 | 17 | N | --- | 91 | --- |
| 2 | Merozoite | 1 | 10 | 5 | 1 | N | --- | 16 | 16 |
| 3 | Schizont | 1 | 11 | 17 | 4 | Y | 5. 0. 3 | 32 | 227 |
| 3 | Schizont | 2 | 11 | 17 | 2 | Y | 4. 0. 2 | 30 | --- |
| 3 | Schizont | 3 | 14 | 23 | 3 | Y | 5. 0. 1 | 40 | --- |
| 3 | Schizont | 4 | 13 | 15 | 0 | Y | 4. 2. 0 | 28 | --- |
| 3 | Schizont | 5 | 9 | 12 | 2 | Y | 5. 0. 2 | 23 | --- |
| 3 | Schizont | 6 | 15 | 17 | 0 | Y | 7. 2. 0 | 32 | --- |
| 3 | Schizont | 7 | 13 | 7 | 0 | Y | 5. 1. 0 | 20 | --- |
| 3 | Schizont | 8 | 10 | 11 | 1 | Y | 5. 1. 0 | 22 | --- |
| 4 | Troph | 1 | 48 | 20 | 13 | N | --- | 81 | 81 |
| 5 | Schizont | 1 | 22 | 11 | 2 | Y | 4. 0. 1 | 35 | 63 |
| 5 | Schizont | 2 | 12 | 15 | 1 | Y | 6. 0. 0 | 28 | --- |
| 6 | Troph | 1 | 40 | 32 | 15 | Y | 2. 1. 0 | 87 | 148 |
| 6 | Troph | 2 | 29 | 18 | 14 | Y | 1. 0. 0 | 61 | --- |
| 7 | Troph | 1 | 63 | 16 | 0 | N | --- | 79 | 79 |
| 8 | Schizont | 1 | 22 | 10 | 5 | N | --- | 37 | 120 |
| 8 | Schizont | 2 | 23 | 8 | 0 | N | --- | 31 | --- |
| 8 | Schizont | 3 | 18 | 6 | 4 | N | --- | 28 | --- |
| 8 | Schizont | 4 | 11 | 13 | | N | --- | 24 | --- |
| Totals: | | | 447 | 355 | 101 | | | 903 | 903 |

**Experiment** 20220722 Nup221 M2 20hpi

| Cell number | Cell Stage | Nucleus number | Number of GFP Nups | Number of Myc nups | Number of colocalized Nups | CP (Y or N) | Number of nups around CP (GFP, Myc, colocalized) | Total number of nups in nucleus | Total number nups in cell |
|---|---|---|---|---|---|---|---|---|---|
| 1 | Schizont | 1 | 29 | 37 | 2 | Y | 2, 2, 1, | 68 | 288 |
| 1 | Schizont | 2 | 28 | 52 | 5 | Y | 2. 2. 1. | 85 | --- |
| 1 | Schizont | 3 | 17 | 46 | 5 | Y | 0. 5. 1. | 68 | --- |
| 1 | Schizont | 4 | 16 | 46 | 5 | Y | 2. 3. 0 | 67 | --- |
| 2 | Schizont | 1 | 23 | 23 | 2 | Y | 0. 2. 0 | 48 | 269 |
| 2 | Schizont | 2 | 17 | 23 | 3 | Y | 6. 0. 2 : 1. 5. 0 | 43 | --- |
| 2 | Schizont | 3 | 13 | 28 | 7 | Y | 2. 0. 3 | 48 | --- |
| 2 | Schizont | 4 | 14 | 20 | 4 | Y | 1. 1. 0 : 1. 1. 2 | 38 | --- |
| 2 | Schizont | 5 | 24 | 22 | 1 | Y | 0. 6. 0 : 2. 3. 0 | 47 | --- |
| 2 | Schizont | 6 | 22 | 20 | 3 | Y | 0. 2. 1 : 3. 1. 0 | 45 | --- |
| 3 | Merozoite | 1 | 15 | 28 | 5 | N | --- | 48 | 48 |
| 4 | Schizont | 1 | 13 | 42 | 8 | Y | 0. 1. 0 : 0. 7. 1 | 63 | 275 |
| 4 | Schizont | 2 | 22 | 43 | 5 | Y | 1. 3. 1 | 70 | --- |
| 4 | Schizont | 3 | 12 | 16 | 1 | Y | 1. 1. 0 | 29 | --- |
| 4 | Schizont | 4 | 12 | 26 | 1 | Y | 0. 2. 1 | 39 | --- |
| 4 | Schizont | 5 | 16 | 31 | 3 | Y | 0. 1. 0 : 0. 3. 0 | 50 | --- |
| 4 | Schizont | 6 | 4 | 17 | 3 | Y | 1. 1. 2 | 24 | --- |
| 5 | Schizont | | | | | N | --- | 0 | --- |
| 5 | Schizont | | | | | N | --- | 0 | --- |
| 6 | Schizont | 1 | 26 | 23 | 7 | Y | 5. 2. 0 | 56 | 295 |
| 6 | Schizont | 2 | 32 | 19 | 5 | Y | 5. 2. 1 | 56 | --- |
| 6 | Schizont | 3 | 40 | 36 | 9 | Y | 4. 1. 1 : 4. 0. 1 | 85 | --- |
| 6 | Schizont | 4 | 19 | 16 | 7 | Y | 5. 1. 0 | 42 | --- |
| 6 | Schizont | 5 | 27 | 23 | 6 | Y | 4. 1. 0 | 56 | --- |
| 7 | Troph | 1 | 25 | 35 | 16 | N | --- | 76 | 76 |
| 8 | Troph | | | | | N | --- | 0 | --- |
| 9 | Troph | 1 | 38 | 48 | 15 | N | --- | 101 | 101 |
| 10 | Schizont | 1 | 8 | 18 | 0 | N | --- | 26 | 197 |
| 10 | Schizont | 2 | 16 | 49 | 3 | Y | 1. 2. 1 | 68 | --- |
| 10 | Schizont | 3 | 14 | 39 | 2 | Y | 0. 6. 0 | 55 | --- |
| 10 | Schizont | 4 | 12 | 34 | 2 | Y | 1. 0. 0 | 48 | --- |
| Totals: | | | 554 | 860 | 135 | 0 | 0 | 1549 | 1549 |

**Experiment** 20221008 Nup221 M1 10hpi

| Cell number | Cell Stage | Nucleus number | Number of GFP Nups | Number of Myc nups | Number of colocalized Nups | CP (Y or N) | Number of nups around CP (GFP, Myc, colocalized) | Total number of nups in nucleus | Total number nups in cell |
|---|---|---|---|---|---|---|---|---|---|
| 1 | Merozoite | 1 | 14 | 11 | 0 | Y | 5, 0, 0 | 25 | 201 |
| 1 | Merozoite | 2 | 19 | 10 | 0 | Y | 5. 1. 0 | 29 | --- |
| 1 | Merozoite | 3 | 18 | 11 | 2 | Y | 4. 0. 2 | 31 | --- |
| 1 | Merozoite | 4 | 16 | 9 | 2 | Y | 4. 1. 1 | 27 | --- |
| 1 | Merozoite | 5 | 14 | 11 | 1 | N | --- | 26 | --- |
| 1 | Merozoite | 6 | 12 | 6 | 4 | Y | 4. 0. 2 | 22 | --- |
| 1 | Merozoite | 7 | 16 | 6 | 0 | N | --- | 22 | --- |
| 1 | Merozoite | 8 | 10 | 7 | 2 | N | --- | 19 | --- |
| 2 | Troph | 1 | 95 | 68 | 75 | Y | 0. 2. 6 : 2. 1. 1 | 238 | 238 |
| 3 | Merozoite | 1 | 18 | 9 | 6 | N | --- | 33 | 273 |
| 3 | Merozoite | 2 | 18 | 8 | 6 | N | --- | 32 | --- |
| 3 | Merozoite | 3 | 25 | 7 | 8 | N | --- | 40 | --- |
| 3 | Merozoite | 4 | 21 | 7 | 4 | N | --- | 32 | --- |
| 3 | Merozoite | 5 | 15 | 5 | 6 | N | --- | 26 | --- |
| 3 | Merozoite | 6 | 12 | 9 | 9 | N | --- | 30 | --- |
| 3 | Merozoite | 7 | 28 | 6 | 3 | N | --- | 37 | --- |
| 3 | Merozoite | 8 | 31 | 7 | 5 | N | --- | 43 | --- |
| 4 | Schizont | 1 | 17 | 11 | 16 | Y | 0. 2. 5 : 1. 3. 3 | 44 | 179 |
| 4 | Schizont | 2 | 12 | 26 | 25 | Y | 0. 3. 4 : 0. 1.5 : 0. 4. 2 | 63 | --- |
| 4 | Schizont | 3 | 13 | 12 | 6 | Y | 1. 2. 4 | 31 | --- |
| 4 | Schizont | 4 | 10 | 15 | 16 | Y | 0. 4. 4 | 41 | --- |
| 6 | Schizont | 1 | 12 | 31 | 22 | Y | 1. 4. 3 | 65 | 65 |
| Totals: | | | 446 | 292 | 218 | | | 956 | 956 |

**Experiment** 20221008 Nup221 M1 20hpi

| Cell number | Cell Stage | Nucleus number | Number of GFP Nups | Number of Myc nups | Number of colocalized Nups | CP (Y or N) | Number of nups around CP (GFP, Myc, colocalized) | Total number of nups in nucleus | Total number nups in cell |
|---|---|---|---|---|---|---|---|---|---|
| 3 | Troph | 1 | 83 | 109 | 52 | Y | 0, 3, 3 : 2, 2, 3 : 0, 4, 1 : 1, 2, 2 | 244 | 244 |
| 4 | Troph | 1 | 51 | 38 | 15 | N | --- | 104 | 160 |
| | Troph | 2 | 12 | 35 | 9 | Y | 0. 2. 0 : 0. 1. 0 | 56 | --- |
| Totals: | | | 146 | 182 | 76 | | | 404 | 404 |
